# Brain Connectivity Correlates of Breathing and Cardiac Irregularities in SUDEP: A Resting-State fMRI Study

**DOI:** 10.1101/2023.05.19.541412

**Authors:** Michalis Kassinopoulos, Nicolo Rolandi, Laren Alphan, Ronald M. Harper, Joana Oliveira, Catherine Scott, Lajos R. Kozák, Maxime Guye, Louis Lemieux, Beate Diehl

## Abstract

Sudden unexpected death in epilepsy (SUDEP) is the leading cause of premature mortality among people with epilepsy. Evidence from witnessed and monitored SUDEP cases indicate seizure-induced cardiovascular and respiratory failures; yet, the underlying mechanisms remain obscure. SUDEP occurs often during the night and early morning hours, suggesting that sleep or circadian rhythm-induced changes in physiology contribute to the fatal event. Resting-state fMRI studies have found altered functional connectivity between brain structures involved in cardiorespiratory regulation in later SUDEP cases and in individuals at high-risk of SUDEP. However, those connectivity findings have not been related to changes in cardiovascular or respiratory patterns. Here, we compared fMRI patterns of brain connectivity associated with regular and irregular cardiorespiratory rhythms in SUDEP cases with those of living epilepsy patients of varying SUDEP risk, and healthy controls. We analysed resting-state fMRI data from 98 patients with epilepsy (9 who subsequently succumbed to SUDEP, 43 categorized as low SUDEP risk (no tonic-clonic seizures (TCS) in the year preceding the fMRI scan), and 46 as high SUDEP risk (>3 TCS in the year preceding the scan)) and 25 healthy controls. The global signal amplitude (GSA), defined as the moving standard deviation of the fMRI global signal, was used to identify periods with regular (‘low state’) and irregular (‘high state’) cardiorespiratory rhythms. Correlation maps were derived from seeds in twelve regions with a key role in autonomic or respiratory regulation, for the low and high states. Following principal component analysis, component weights were compared between the groups. We found widespread alterations in connectivity of precuneus/posterior cingulate cortex in epilepsy compared to controls, in the low state (regular cardiorespiratory activity). In the low state, and to a lesser degree in the high state, reduced anterior insula connectivity (mainly with anterior and posterior cingulate cortex) in epilepsy appeared, relative to healthy controls. For SUDEP cases, the insula connectivity differences were inversely related to the interval between the fMRI scan and death. The findings suggest that anterior insula connectivity measures may provide a biomarker of SUDEP risk. The neural correlates of autonomic brain structures associated with different cardiorespiratory rhythms may shed light on the mechanisms underlying terminal apnea observed in SUDEP.

## 1. Introduction

Sudden unexpected death in epilepsy (SUDEP) is the leading cause of premature death in patients with intractable epilepsy, with an annual incidence estimated at approximately 1.2 per 1,000 persons with epilepsy (Keller et al., 2018; Sveinsson et al., 2017; Thurman et al., 2017). SUDEP is defined as ‘the sudden, unexpected, witnessed or unwitnessed, nontraumatic and nondrowning death in patients with epilepsy, with or without evidence for a seizure and excluding documented status epilepticus, in which postmortem examination does not reveal a structural or toxicologic cause for death’ (Nashef et al., 2012). SUDEP imposes a substantial public health burden (Thurman et al., 2017) and, among neurological disorders, ranks second only to stroke in terms of potential years of life lost (Thurman et al., 2014). Frequent generalized and focal-to-bilateral tonic-clonic seizures (TCS; Fisher et al., 2017) are the greatest risk factors. Sleep and nocturnal TCS appear to facilitate SUDEP (Ali et al., 2017; Ryvlin et al., 2013). The pathophysiology of SUDEP remains poorly understood (Devinsky et al., 2016; Sveinsson et al., 2020), and seizure control is considered the most effective strategy for reducing risk of the fatal outcome. However, currently, strategies for assessing SUDEP risk on an individual basis are still lacking.

Epilepsy monitoring unit data from the MORTEMUS study (MORTality in Epilepsy Monitoring Unit Study) suggest that SUDEP results from cardiorespiratory dysfunction induced by TCS (Ryvlin et al., 2013). Vilella et al. (2019) found that post-ictal central apnea occurs in one out of five TCS and was present in near-SUDEP and SUDEP cases, suggesting that cessation of breathing may represent an important SUDEP biomarker. Cardiovascular parameters may also provide markers; heart rate and its variability, measured in the peri-ictal period (*e.g.* post-ictal mean heart rate; Arbune et al., 2020), are associated with markers of seizure severity that have been linked to SUDEP such as the presence of TCS and the duration of post-ictal generalized EEG suppression (PGES). However, although some patients succumb to SUDEP after a few seizures, others survive hundreds of similar attacks, which suggests the presence of additional pathophysiological mechanisms in SUDEP victims (Devinsky and Sisodiya, 2020). Intriguingly, reduced interictal heart rate variability (HRV) measured during wakefulness has also been associated with SUDEP (DeGiorgio et al., 2010; Sivathamboo et al., 2021), raising the possibility of chronic impairment in autonomic regulation in SUDEP. Further evidence of the role of chronic dysregulation is provided by the recent observation of abnormal heart rate responses during and after hyperventilation in patients who subsequently died of SUDEP (Szurhaj et al., 2021), and of volume changes in brain regions with key roles in autonomic regulation (Allen et al., 2019b; Mueller et al., 2018).

Functional MRI (fMRI) is a non-invasive neuroimaging tool that can evaluate functional connectivity (FC) between brain structures at a whole-brain level. Early fMRI studies focusing on FC revealed altered connectivity of regions involved in cardiorespiratory regulation such as the anterior cingulate cortex, thalamus and regions of the brainstem in SUDEP cases and patients at high risk of SUDEP (Allen et al., 2019a, 2017; Tang et al., 2014). These studies, however, did not take into consideration the time-varying nature of FC observed on the scale of seconds to minutes, which is believed to provide a more holistic understanding of the brain functional organization (Chang and Glover, 2010; Preti et al., 2017). There is accumulating evidence that FC dynamics change with different autonomic and sleep states (Chang et al., 2013; Haimovici et al., 2017). Thus, examining the changes in FC occurring in different dynamic patterns of breathing or cardiovascular action, such as those that appear during different sleep states or other provocations, may reveal new insights into mechanisms that contribute to SUDEP different from those FC values obtained in static physiological conditions.

In addition to the apparent role of sleep and nocturnal TCS, recent observations suggest that more attention on the state-dependent nature of autonomic manifestations is warranted. A recent study suggests a stronger association between SUDEP and post-ictal rather than ictal central apnea (Vilella et al., 2019); whereas, another study found abnormally low HRV in SUDEP during wakefulness, but not during sleep (Sivathamboo et al., 2021). Thus, the examination of the state-dependent nature of autonomic manifestations using resting-state fMRI seems promising. In particular, the observation that subjects that exhibit strong fluctuations in heart rate and breathing patterns during an fMRI scan are also characterized by an elevated global signal amplitude (GSA), which refers to strong BOLD fMRI fluctuations globally in the brain (Orban et al., 2020; Power et al., 2017; Xifra-Porxas et al., 2021), may be a key to better understanding the physiology of SUDEP.

In this work, we sought to characterize the patterns of FC in patients who eventually succumbed to SUDEP, living patients of varying SUDEP risk levels, and healthy controls, with regards to variations in the regularity of cardiorespiratory rhythms. First, we use a publicly available fMRI dataset (Van Essen et al., 2013) with concurrent physiological recordings to demonstrate that the association of GSA with cardiac and breathing rhythms holds even within short fMRI scans (∼15 min), with periods of high GSA corresponding to times with irregularities in cardiac or breathing activity such as periods with transient apnea. Second, we characterize patterns of FC in SUDEP cases and epilepsy patients alive at the time of this analysis by employing a state-dependent framework and GSA as a surrogate of the state of autonomic activity. Moreover, given the well-documented success of FC measures in predicting symptom severity in individuals for a range of disorders (Du et al., 2017; Uddin et al., 2013; Yoo et al., 2018), we also investigate whether FC measures in SUDEP cases are associated with the interval between the fMRI scan and SUDEP occurrence.

## 2. Materials and methods

The principal aim was to study the link between brain connectivity and autonomic activity in patients with epilepsy using a large resting-state fMRI dataset that did not comprise physiological recordings. This study consists of two experiments: In the first experiment (Experiment 1), we employed a set of resting-state fMRI data that included concurrent recordings from a photoplethysmograph (PPG) and a respiratory belt, to demonstrate that global signal amplitude (GSA) fluctuations reflect changes in cardiorespiratory activity. Previous studies have defined GSA as the standard deviation of the global signal across the entire scan (Wong et al., 2016, 2013) and showed that GSA is linked to physiological parameters (Orban et al., 2020). Here, we compute GSA over significantly shorter durations, using the sliding window approach, in order to illustrate that the relationship of GSA with physiological parameters also holds at shorter timescales. Experiment 2 consisted of the characterization of the patterns of FC in patients with epilepsy using GSA as a marker of breathing and cardiac irregularities.

### 2.1 Experiment 1: Association of fMRI global signal amplitude with variations in cardiorespiratory rhythms (HCP data)

To demonstrate that the global signal amplitude is linked to cardiorespiratory activity, we examined resting-state fMRI data from the Human Connectome Project (HCP; Van Essen et al., 2013) that included concurrent recordings from a photoplethysmograph (PPG) and a respiratory belt. A description of the preprocessing pipeline for the HCP dataset can be found in the Supplementary Material. Data from a subset of 400 healthy young participants that was characterized by good-quality physiological recordings were included. The global signal, defined as the mean fMRI time-series averaged across all voxels in the grey matter, was computed from the fMRI data after volume realignment and high-pass filtering (0.008 Hz). Subsequently, the global signal amplitude (GSA), defined as the standard deviation of the global signal, was computed in a sliding window manner for window lengths ranging from 10 to 120 sec (or equivalently, 14 to 167 time points) using the Matlab function *movstd*. A one-sample shift was applied between consecutive windows, and the standard deviation computed within a window was assigned to the center of the window in terms of time. In addition, the following four variables were obtained from the physiological recordings: (1) breathing rate; (2) respiration volume as defined in Chang et al. (Chang et al., 2009; i.e. moving standard deviation of respiratory signal with a window length of six seconds); (3) heart rate; and (4) PPG amplitude defined as the amplitude of the oscillatory signal in the PPG (Kassinopoulos and Mitsis, 2021). Subsequently, the moving standard deviation of the physiological variables was also estimated for window lengths ranging from 10 to 120 sec. Then, for each window length, the correlation of the GSA with each of the four physiological variables was computed and averaged across all individuals, in order to find the length that maximizes the correlation without sacrificing temporal resolution.

### 2.2 Experiment 2: Characterization of the GSA-related patterns of FC in patients with epilepsy

#### 2.2.1 Subjects

We retrospectively ascertained cases of SUDEP, and high- and low-risk patients from the University College London Hospitals (UCLH) clinical database that had undergone an EEG-fMRI scan in the period between 2005 and 2015 (Coan et al., 2016). The inclusion criteria were: (1) the availability of a resting-state EEG-fMRI scan; and (2) a high-resolution T^1^-weighted scan. The exclusion criteria were: (1) large brain lesion or previous neurosurgery (we considered large to be anything similar to or greater than a small area of focal cortical dysplasia (FCD) or hippocampal sclerosis – e.g. tumours, cavernomas); (2) incomplete clinical or imaging data (e.g. abandoned scans); and (3) having died in the following years with a cause of death not related to SUDEP. Only patients alive at the time of writing were considered as low-risk or high-risk epilepsy controls.

Out of a cohort of 189 patients who underwent resting-state EEG-fMRI, 14 deaths were identified in the UCLH clinical database, of which 10 were classified as SUDEP based on their death certificate. One SUDEP case was excluded due to the presence of a large brain lesion. The remaining 9 SUDEP cases (5 females, mean age 26.2 ± 6.2) were classified as either probable or definite SUDEP based on the definitions proposed in Nashef et al. (2012). The 9 examined SUDEP cases were matched with 43 low-risk, 46 high-risk patients and 25 healthy controls based on sex and age at the time of scan. High-risk patients were considered those that experienced more than 3 TCS in the year preceding the scan and low-risk patients were considered those that did not experience TCS. Group demographics and clinical details are shown in Suppl. Table 1. The study was approved by our local Research Ethics Committee and all patients gave written informed consent.

#### 2.2.2 Simultaneous EEG-fMRI acquisition

Scanning was performed at the Epilepsy Society (Chalfont St Peter, Buckinghamshire, UK) on a 3.0 Tesla GE (Signa excite HDX) scanner. A 20-min (400 vol) T_2_*-weighted fMRI scan was collected from each subject except for two patients that were scanned for 10-min instead. The fMRI scan was done using a gradient-echo echo-planar-imaging with the following characteristics: repetition time (TR) = 3000 ms, echo time (TE) = 30 ms, flip angle = 90°, matrix size = 64 x 64, field of view (FOV) = 24 x 24 cm^2^, slice thickness = 2.4 mm with 0.6 mm gap, 44 slices, and voxel size = 3.75 x 3.75 x 3 mm^3^. Subjects were instructed to keep their eyes closed, avoid falling asleep, and not think about anything in particular. A T_1_-weighted image was also acquired using a FSPGR (fast spoiled gradient recalled echo) sequence, with the following parameters: matrix size = 256 x 256, FOV = 24 x 24 cm^2^, slice thickness = 1.5 mm, 150 slices, and voxel size = 0.94 x 0.94 x 1.5 mm^3^.

Scalp EEG signals and an electrocardiogram (ECG) signal were simultaneously acquired during fMRI scanning at 5 kHz using a 64-channel MR-compatible EEG system with a cap comprising ring Ag/AgCl electrodes (BrainAmp MR+; Brain Products GmbH, Gilching, Germany). The electrodes were placed according to the 10/20 system and referenced to electrode FCz.

#### 2.2.3 Preprocessing of fMRI data

The preprocessing of fMRI data was conducted using the Statistical Parametric Mapping software (SPM12, Welcome Trust Centre for Neuroimaging, London, UK, http://www.fil.ion.ucl.ac.uk/spm; Friston et al., 2007) in a Matlab environment (R2020a; Mathworks, Natic, Massachusetts, USA). The first five functional volumes were discarded to allow steady-state magnetization to be established and the remaining volumes were realigned to correct for head movements. The structural image of each subject was co-registered to the mean realigned functional volume and, subsequently, underwent tissue segmentation into grey matter, white matter and cerebrospinal fluid tissue compartments. The functional images, as well as the coregistered structural images and tissue compartment masks, were spatially normalized to the Montreal Neurological Institute (MNI) reference space using non-linear transformation.

To account for anatomical variability across participants and reduce thermal noise, all individual functional volumes were smoothed using a 5 mm full-width half-maximum (FWHM) Gaussian kernel. The fMRI time-series were high-passed at 0.008 Hz to avoid spurious correlations that arise from low-frequency fluctuations (Leonardi and Van De Ville, 2015).

We used the frame-wise displacement (FD) introduced by Salek-Haddadi et al. (2006), implemented here as calculated in Power et al. (2012), to identify and exclude subjects with high levels of motion as motion has been shown to lead to systematic biases in FC studies (Kassinopoulos and Mitsis, 2022; Power et al., 2015; Savva et al., 2020; Xifra-Porxas et al., 2021). FD is calculated from the six motion realignment parameters, and reflects the extent of scan-to-scan head motion at each time point. Subjects that were characterized by mean FD larger than 0.25 mm were excluded. For the datasets included in this study, time points with FD larger than 0.2 mm were considered outliers and corrected by linear interpolation.

Finally, to further mitigate the effects of motion and reduce the effects of physiological processes and scanner artifacts, we regressed out the following nuisance effects from all voxel time-series: the first 10 principal components extracted from all white matter voxel time-series (Behzadi et al., 2007), 6 regressors related to cardiac pulsatility artifacts obtained with the convolution model proposed in Kassinopoulos & Mitsis (2021) in conjunction with the R-waves detected in the ECG, and the mean fMRI time-series averaged across all voxels within the grey matter (Macey et al., 2004). The mean grey matter signal was considered a nuisance regressor, as there is accumulating evidence that it reflects changes in heart rate and breathing patterns (Kassinopoulos and Mitsis, 2019; Xifra-Porxas et al., 2021). Note that although GSA, which is used in this study to probe regularities in cardiorespiratory activity, is derived from the grey matter signal, the two aforementioned signals are weakly correlated, with a group average correlation of -0.05 (± 0.25) (for a GSA estimation with a sliding window length of 80 sec).

#### 2.2.4 Seed-based connectivity and spatial distribution pattern analysis

A seed-based FC analysis was employed with seeds consisting of regions with a key role in autonomic and respiratory regulation (Benarroch, 1993; Critchley et al., 2003; Lacuey et al., 2018; Song et al., 2018; Valenza et al., 2019). Specifically, the seeds consisted of the mean time-series of voxels within the following 12 regions: anterior cingulate cortex, anterior insula, posterior insula, thalamus, amygdala, hippocampus, parahippocampal gyrus, precuneus/posterior cingulate cortex (PCu/PCC), cuneus, caudate, putamen and Brodmann area 25. The aforementioned regions were defined based on parcels (i.e. non-overlapping contiguous regions) from the Brainnetome atlas (Fan et al., 2016).

As in Exp. 1, the global signal used to derive GSA was computed as the mean time-series averaged across all voxels in the grey matter, after volume realignment and high-pass filtering (0.008 Hz), and before regressing out nuisance regressors. A window length of 80 sec was used to derive the GSA, as this length was found in Exp. 1 to yield a strong association between the GSA and the physiological variables, while also preserving an adequate temporal resolution (see Results, Section 3.1). Time points that corresponded to the lowest and highest quartile of the GSA trace were assigned to the low and high state, where the low and high states indicate, respectively, times with regular (stable) and irregular (unstable) cardiorespiratory rhythms (Fig. 1A). Time points that corresponded to the second and third quartile were discarded from the analysis.

**Fig. 1.**
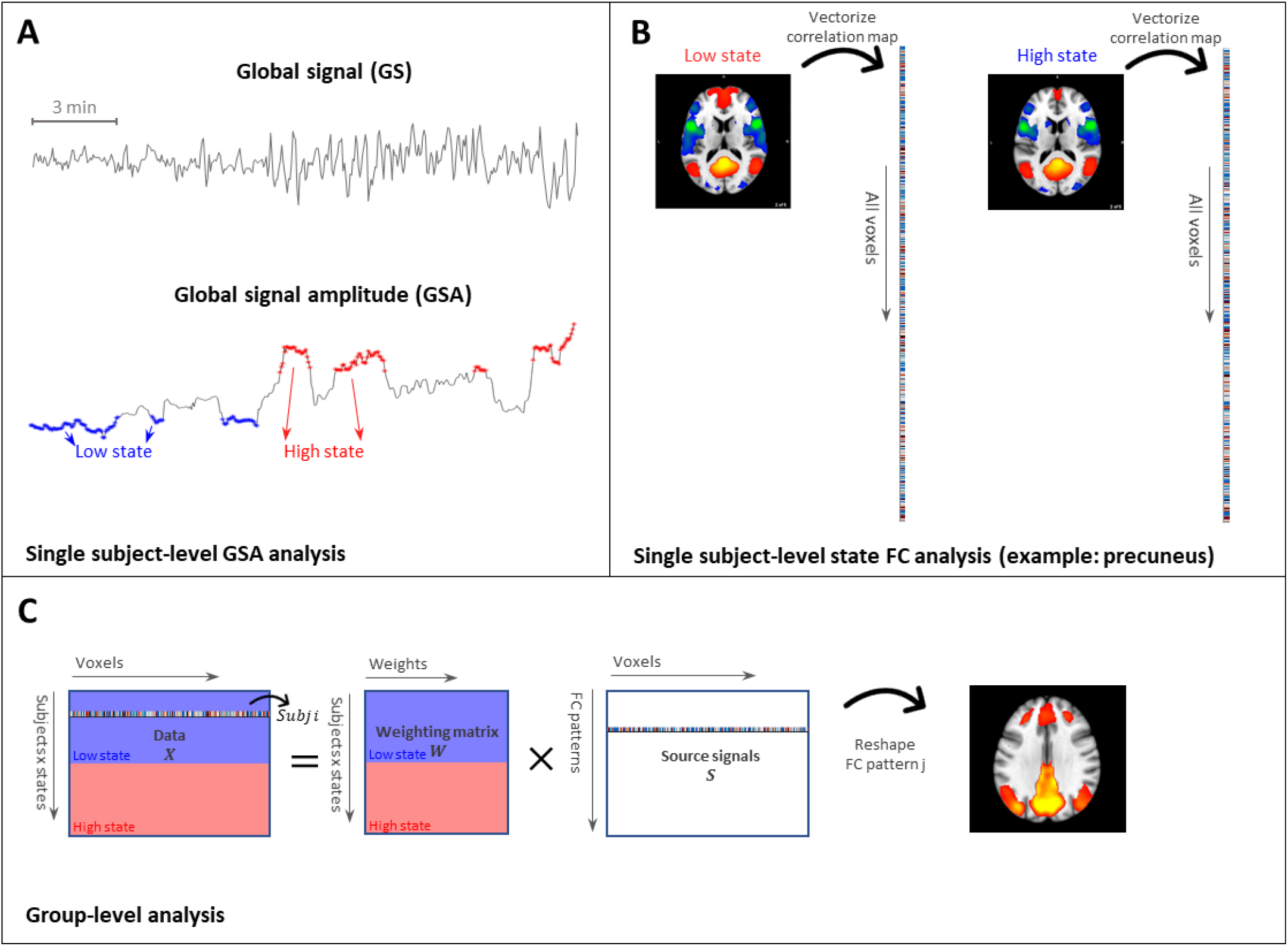
Experiment 2. Analytical framework for characterizing functional connectivity (FC) in individuals based on a linear combination of group-level FC patterns. **(A)** The global signal amplitude (GSA), defined as the moving standard deviation of the fMRI global signal (window length: 80 sec), was computed and used to determine times with regular (‘low state’) and irregular (‘high state’) cardiorespiratory activity. **(B)** Subsequently, for each subject, state and seed region of interest (here, precuneus/posterior cingulate cortex) we computed whole-brain correlation maps represented in vectorized form. **(C)** Finally, for each seed, the connectivity strength vectors were concatenated across subjects and states, and the resulting two-dimensional matrix (left of the equal sign) was fed into a principal component analysis (PCA) to derive the underlying group-level FC patterns.

The fMRI time-series derived for each seed was correlated with all regions in the grey matter on a voxel-wise basis (Pearson correlation), considering the low and high states, separately. Time points that corresponded to the low state were concatenated before computing the Pearson correlation between seed and voxel timeseries. Likewise, time points in the high state were concatenated before computing the Pearson correlation. The correlation values were then converted into z-scores using Fisher’s transform (Fig. 1B). For each seed, the correlation maps were vectorized and concatenated across subjects and states, resulting in a two-dimensional matrix where rows represent subjects and states (e.g. a row may correspond to subject *j* in the low state), and columns correspond to voxels. Finally, the resulting matrix was decomposed through principal component analysis (PCA) to generate a set of FC patterns, also known as eigenconnectivities (Leonardi et al., 2013), and a set of weights reflecting the degree to which a subject expresses each of the FC patterns in a given state (Fig. 1C). To reduce the number of tests, the subsequent analysis was restricted to the components explaining the highest fraction of variance with a 90% of cumulative variance.

#### 2.2.5 Permutation-based statistical analysis

The PCA weights obtained for each component were compared between the four groups (SUDEP, high-risk patients, low-risk patients, controls) using ANOVA, considering the weights associated to the low and high states separately. The *F*-statistics associated with pairs of components and states were mapped to *p*-values based on a null distribution generated using permutation tests. To generate the null distribution of *F*-statistics, 10,000 permutations were performed where in each permutation the subjects were randomly assigned to one of the four groups keeping the size of each group same to the size of the real groups. Subsequently, the *F*-statistics obtained from all examined components (i.e. the M most-significant components that corresponded to a 90% cumulative variance), the 12 seeds and the two states were pooled to generate the null distribution. The alpha level was set at *p* < 0.05 which was corrected for multiple comparisons (M=150, twelve seeds, two states) using Bonferroni correction. For the components found to discriminate between the four groups, we explored the possibility that the involvement of a component in brain connectivity is associated with the time interval between the fMRI scan and occurrence of SUDEP using the Pearson’s correlation coefficient.

#### 2.2.6 Large-scale network involvement

Finally, we estimated the level of involvement of the seven large-scale networks reported by Yeo et al. (2011) for each connectivity component found to discriminate between the four groups. This was done by calculating each component’s spatial overlap (Sørensen-Dice coefficient) with Yeo’s large scale networks using the *ICN_Atlas* toolbox (Kozák et al., 2017). The sign of the connectivity component was taken as that of its Sørensen-Dice coefficient.

## 3. Results

### 3.1 Experiment 1: Association of fMRI global signal amplitude with variations in cardiorespiratory rhythms (HCP data)

We found the GSA to be strongly correlated with the standard deviation of each of the four examined physiological variables (breathing rate, respiration volume, heart rate and PPG amplitude; Suppl. Fig. 1) in the HCP data (Van Essen et al., 2013). The strongest association was observed for respiration volume; furthermore, increasing the window length from 20 to 80 sec led to a significant increase of the correlation from 0.29 (±0.02) to 0.53 (±0.03) which remained at similar levels for longer lengths. A window length of approximately 80 sec was also found to increase the correlation of GSA with breathing rate (0.34±0.03) and PPG amplitude (0.30±0.03). Fig. 2 shows the breathing-related and fMRI signals for a subject where the levels of GSA co-fluctuate with variations in respiration volume. For instance, we observe that periods with strong fluctuations in respiration volume (e.g. 200-500 sec) were characterized by higher levels of GSA as compared to periods with a regular breathing activity (e.g. 500-600 sec). Finally, we note that while a similar association of GSA with respiration volume was observed, for several subjects, high levels of GSA often corresponded to periods with transient apnea (i.e. brief pauses of breathing activity; Suppl. Fig. 2A), transient increase in heart rate (Suppl. Fig. 2B) or variations in PPG amplitude (Suppl. Fig. 2C). Therefore, GSA is strongly driven by variations in cardiovascular and breathing activity, although it cannot distinguish the former from the latter.

**Fig. 2.**
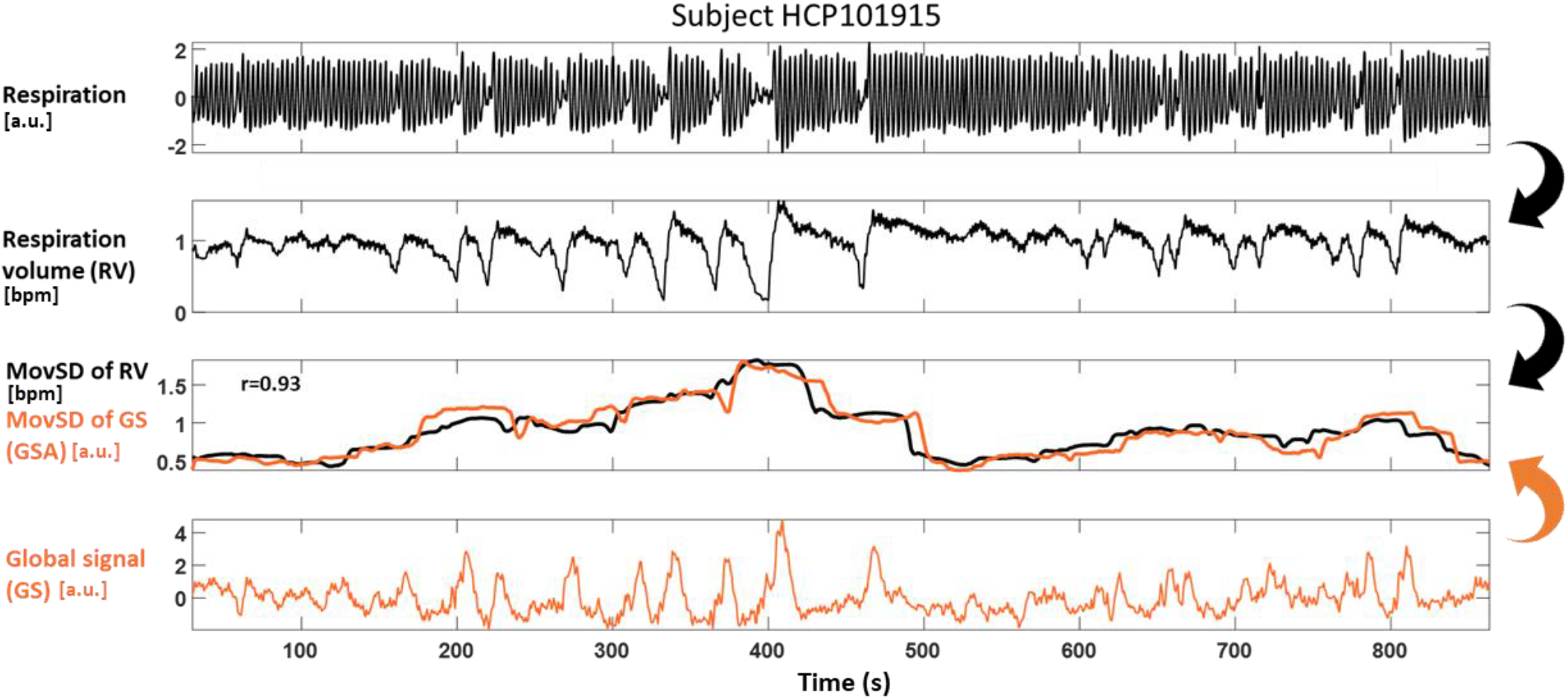
Experiment 1. Illustration of strong correlation (r=0.93) between global signal amplitude (GSA) and irregularity in breathing for an HCP dataset subject. (1^st^ row) Respiration monitored with a respiratory bellows placed around the chest. (2^nd^ row) Respiration volume (RV) defined as the moving standard deviation of respiration using a window length of 6 sec as in Chang et al. (2009). (3^rd^ row) Moving standard deviation of RV (black line) and global signal (GS; orange line) using a window length of 80 sec. (4^th^ row) FMRI GS defined as the mean time-series averaged across all voxels in the grey matter. Note that high values of GSA (e.g. between 200 and 500 sec) indicate periods with strong fluctuations in RV, whereas low values of GSA (e.g. 500-600 sec) indicate periods with regular breathing. Resting-state data from HCP subject HCP101915 (Van Essen et al. 2013). For an illustration of the association between GSA and breathing rate, heart rate and PPG amplitude, see Suppl. Fig. 2.

### 3.2 Experiment 2: Characterization of the GSA-related patterns of FC in patients with epilepsy

Three low-risk and one high-risk epilepsy patients were excluded due to excessive motion (mean FD > 0.25 mm) resulting in a cohort of 119 subjects (57 women; mean age 30.4 ± 8.4): healthy controls: 25; low-risk: 40; high-risk: 45; SUDEP: 9. The sex and age distributions were similar between the four groups (Suppl. Table 1). In addition, the four groups exhibited similar levels of fMRI scan-wise motion as assessed with ANOVA (*F* = 1.3; *p* > 0.05).

#### 3.2.1 Identification of autonomic structures with significant group differences

The seed-based correlation maps of twelve brain structures with a key role in autonomic regulation were computed for all subjects in the low and high state (i.e. considering functional volumes with GSA values in the lowest and highest quartile, respectively). For each of the twelve seeds, 150 principal components were found to explain about 90% of the variance in the correlation maps. Among the twelve seeds, only three (anterior insula, PCu/PCC and cuneus) yielded components for which the weights could discriminate between the four groups (i.e. healthy controls, low-risk, high-risk and SUDEP patients; Fig. 3). More specifically the *F*-statistic, which reflects the degree to which the mean of component weights differed between the four groups, was significant (chance level: *F* < 9.4, corresponding to a *p*-value of 2 × 10^−5^ under a null distribution of 10,000 permutations) for the following components: Anterior insula #3 in the low state (*F* = 13.8, *p* < 10^-7^); PCu/PCC #2 in the high state (*F* = 12.0, *p* < 10^-6^); and Cuneus #1 in the high state (*F* = 10.2, *p* < 10^-5^).

**Fig. 3.**
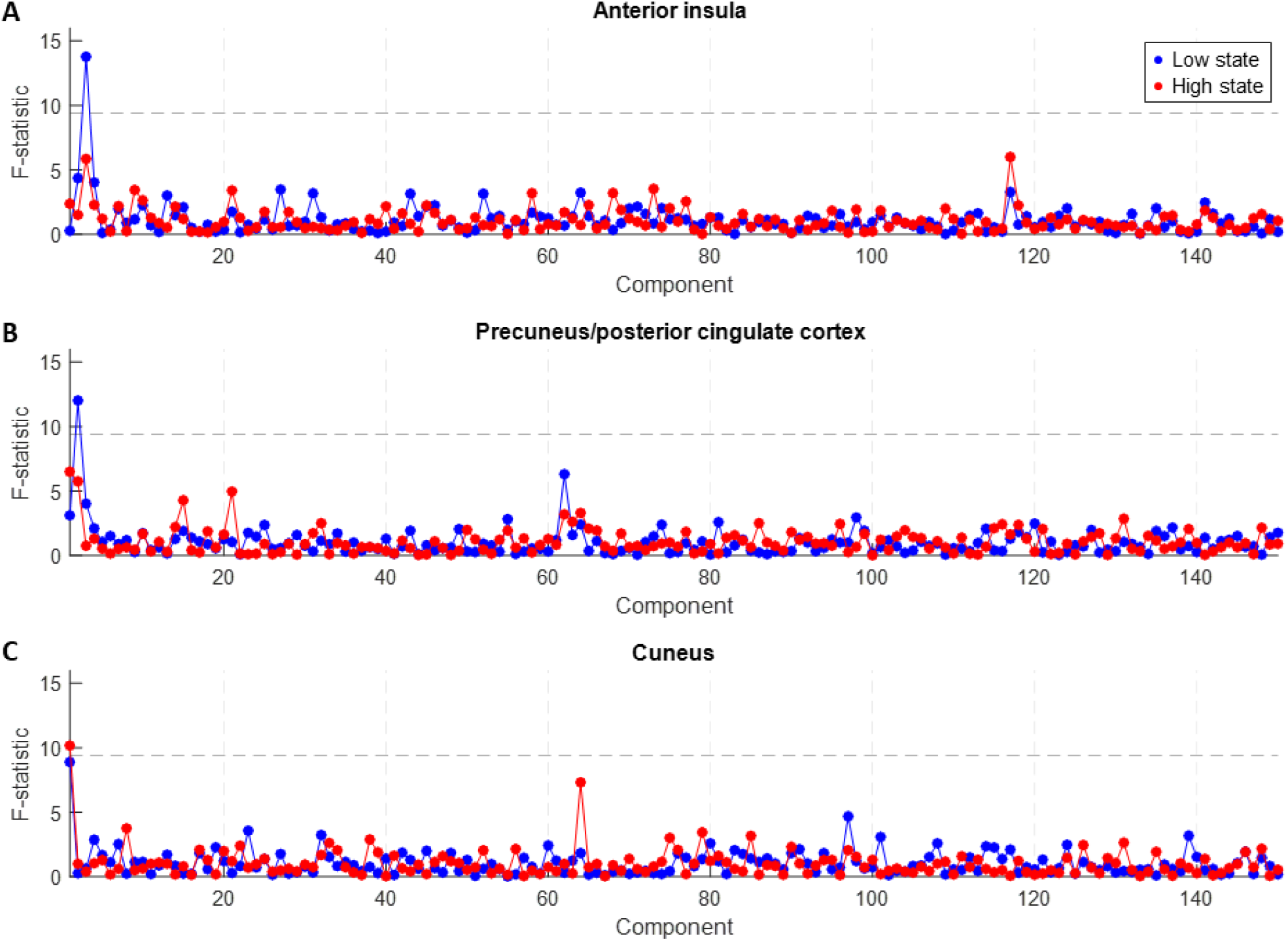
Exp. 2. *F*-statistics for assessing dispersion between groups based on FC patterns of (A) anterior insula, (B) precuneus/posterior cingulate cortex (PCu/PCC) and (C) cuneus. The dashed line indicates the chance level (*p* < 0.05, Bonferroni corrected), as determined by permutation distribution. All three regions had one of the first three principal components with an *F*-statistic at above chance level. None of the other nine seed regions examined here was found to yield a significant component in terms of *F*-statistic.

#### 3.2.2 Functional connectivity (FC) patterns and relationship with interval between the time of fMRI scan acquisition and SUDEP

In this section we examine the FC patterns specific to the 3 structures identified in the previous: Anterior insula, PCu/PCC and Cuneus.

##### Anterior insula seed connectivity

Fig. 4A shows the mean anterior insula seed-based correlation map (averaged across subjects and time) used as a reference for the interpretation of the different component weights observed across the groups. The anterior insula was found to be positively correlated with the anterior cingulate cortex, regions of the middle frontal gyrus, postcentral gyrus, inferior parietal lobe and posterior insula, and negatively correlated with the posterior cingulate cortex, and regions of the middle temporal gyrus and frontal gyrus. The FC pattern of component #3 that was found to be the most discriminant principal component of anterior insula, exhibited positive weights in the posterior cingulate cortex, and regions of the middle temporal gyrus and frontal gyrus, and negative weights in the anterior cingulate cortex, and anterior and posterior insula (Fig. 4B).

**Fig. 4.**
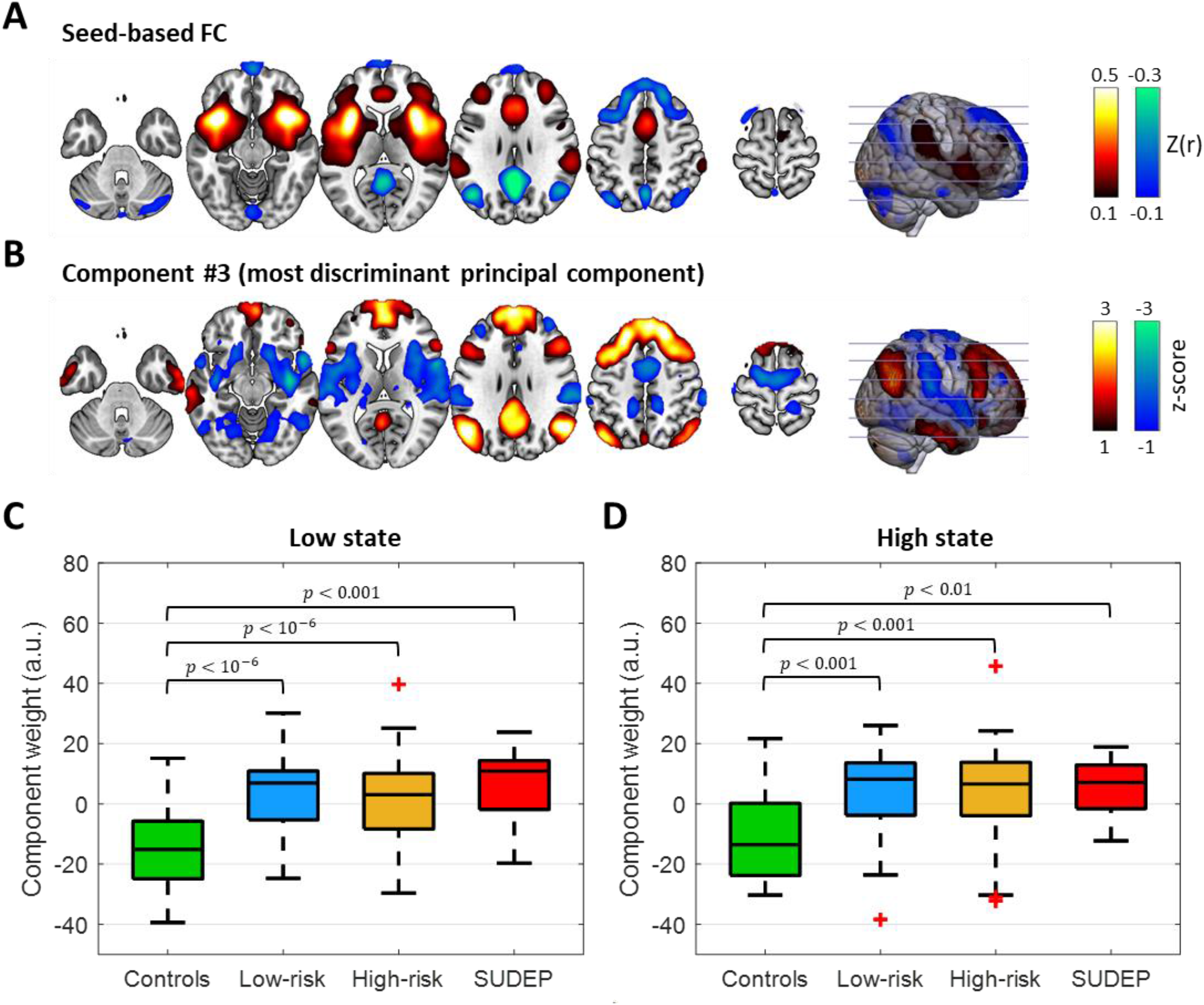
Exp. 2. Involvement of FC pattern #3 of anterior insula connectivity in the low and high (GSA) states. (**A**) Seed-based correlation map with the seed placed in the anterior insula averaged across subjects and time. (**B**) FC pattern of component #3 derived from the anterior insula connectivity profiles of all subjects through PCA. Component weights of the four groups in the (**C**) low and (**D**) high state (the bottom and top of each box correspond to the 25^th^ and 75^th^ percentiles of the sample distribution, the line in the box corresponds to the median and the crosses indicate outliers, defined as values that are more than 1.5 times the interquartile range away from the edges of the box). The FC pattern of component #3 was to a large degree opposite of the anterior insula connectivity profile. Thus, a positive component weight as found in the epilepsy groups can be interpreted as weakening of the anterior insula connectivity with its typical connections (e.g. anterior and posterior cingulate cortex). For the spatial involvement of large-scale networks in FC pattern #3, see Suppl. Fig. 5.

It can be seen that the FC pattern #3 of anterior insula is to a large degree the inverse of the anterior insula seed-based correlation map, and thus a positive weight for the involvement of this pattern on a subject can be interpreted as decline in anterior insula connectivity as compared to the mean connectivity observed in the entire cohort. In other words, both positive and negative correlation values of the anterior insula’s connectivity, with the anterior and posterior cingulate cortices respectively, are weaker. Similarly, a negative weight of FC pattern #3 for within-subject involvement indicates stronger anterior insula connections with respect to the mean connectivity. In both the low and high states, the epilepsy groups were characterized by significantly elevated component weights (*p* < 0.01; Fig. 4C-D). In addition, the majority of individuals with epilepsy were characterized by a positive component weight in contrast to negative component weights for most healthy controls. Furthermore, for the SUDEP cases, the anterior insula FC pattern #3 component weights observed in the high state were found to be strongly anti-correlated with the interval between the fMRI scan and time of SUDEP (*r* = -0.74, *p* = 0.02, uncorrected for the 6 tests performed; Fig. 5), whereas this association was not observed in the low state (*p* > 0.05).

**Fig. 5.**
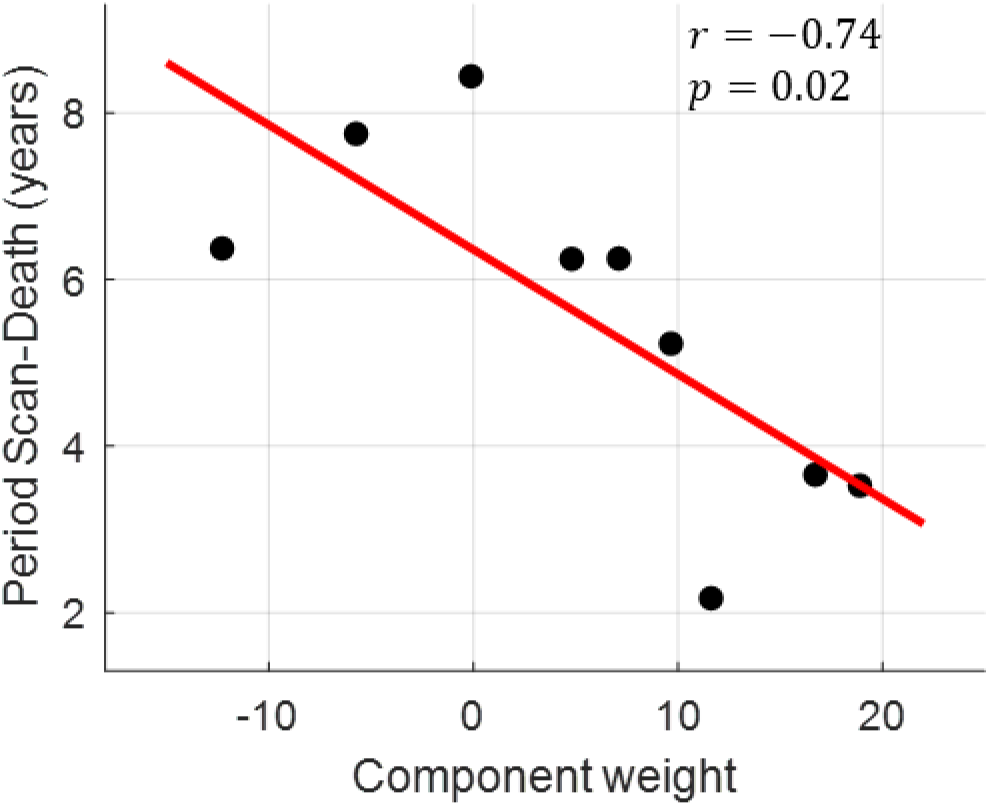
Exp. 2. Strength of anterior insula connectivity linked to the interval between the fMRI scan and time of SUDEP. The weight of component #3 in the high state (irregular cardiorespiratory activity) decreased with interval length (*r* = -0.74, *p* = 0.02). Given that the FC pattern of component #3 (Fig. 4B) is, to a large extent, the inverse of the mean anterior insula connectivity (Fig. 4A), a positive weight for the within-subject involvement can be interpreted as reduced anterior insula connectivity. Thus, the inverse relationship between component weight and scan-SUDEP interval indicates that patients that died relatively soon after the scan exhibited weak anterior insula connectivity.

##### PCu/PCC seed connectivity

The correlation map of the PCu/PCC averaged across subjects and time consisted of positive correlations with the posterior middle and superior temporal gyri, and the medial frontal gyrus, and negative correlations with the precentral and postcentral gyri, inferior frontal gyrus, anterior superior temporal gyrus and anterior insula cortex (Suppl. Fig. 3A). The FC pattern of component #2 (most discriminant principal component of PCu/PCC) consisted of positive weights in the precuneus/posterior cingulate cortex superior occipital cortex, middle temporal gyrus, angular gyrus, and middle and medial frontal gyri (Suppl. Fig. 3B). In addition, it exhibited negative weights in regions close to the cerebral hemispheres and brainstem, such as the amygdala, hippocampus, parahippocampal gyrus, putamen, claustrum, superior temporal gyrus and fusiform gyrus. In the low state, the three epilepsy groups exhibited significantly reduced weights for component #2 (*p* < 0.001; Suppl. Fig. 3C), with the majority of patients exhibiting negative weights and the majority of controls positive weights. Similar observations were made in the high state (Suppl. Fig. 3D). The weights of PCu/PCC component #2 did not show any association with the interval between the fMRI scan and the occurrence of SUDEP.

##### Cuneus seed connectivity

The cuneus correlation map averaged across subjects and time presented negative correlations with the inferior parietal lobe, inferior frontal gyrus and inferior temporal gyrus (Suppl. Fig. 4A). The FC pattern of component #1 (most discriminant component of cuneus) consisted of positive weights in the ventral anterior cingulate cortex, precentral and postcentral gyrus, superior temporal gyrus as well as cuneus, and negative weights in the posterior cingulate cortex, regions of the superior and middle temporal gyrus, thalamus, and frontal gyrus (Suppl. Fig. 4B). In the low state, the three epilepsy groups exhibited significantly higher component weights than healthy controls (*p* < 0.05; Suppl. Fig. 4C). The higher component weights of low-risk and high-risk patients as compared to controls were also observed in the high state (Suppl. Fig. 4D). The weights of cuneus component #1 did not show any association with the interval between the fMRI scan and the occurrence of SUDEP.

##### Large-scale network involvement

With regards to the spatial involvement of the Yeo et al. (2011) large-scale networks in the connectivity patterns revealed in this work, the anterior insula component #3 was found to reflect engagement of the anterior insula with regions of the default mode network and, to a lesser degree, frontoparietal network, and disengagement with the ventral attention and somatomotor network (Suppl. Fig. 5). The PCu/PCC component #2 was found to engage connectivity with regions that partly belong to the default mode network. Although widespread regions of this component were found to be disengaged with the PCu/PCC, they did not resemble any of the large-scale networks. Finally, the cuneus component #1 was found to reflect mainly engagement of the cuneus with regions of the somatomotor network, and disengagement with regions of the default mode network.

## 4. Discussion

This study utilized resting-state fMRI data to examine FC in patients with drug-resistant epilepsy who subsequently died of SUDEP. Given the strong link between time-varying FC and variations in autonomic activity (Chang et al., 2013; Mulcahy et al., 2019; Nikolaou et al., 2016), particular attention was placed on characterizing connectivity in periods with regular and irregular cardiorespiratory activity, separately. We also wanted to examine whether any of the observed effects is linked to the risk for SUDEP.

We first demonstrated that the moving standard deviation of fMRI global signal, termed here global signal amplitude (GSA), is elevated at periods with strong fluctuations in breathing and cardiac activity (Experiment 1) using data in the public domain. Specifically, we showed that GSA is elevated during periods of the order of 1 minute that present at least one of the following features: a transient apnea, variations in respiratory volume, transient increases in heart rate or variations in the amplitude of the PPG (Suppl. Fig. 1); In this work, such periods correspond to a ‘high GSA state’. Our study extends previous findings by demonstrating that the association of GSA and variations in cardiorespiratory rhythms previously found across subjects and fMRI runs (Kassinopoulos and Mitsis, 2019; Orban et al., 2020; Power et al., 2017), is also present within a run at the scale of minutes. This finding indicates that previously-collected fMRI datasets that did not include monitoring of physiological processes could be revisited to study aspects of the autonomic nervous system. This outcome is particularly relevant for fMRI studies examining rare diseases and phenomena with a low incidence rate such as SUDEP.

Subsequently, we characterized FC in 9 SUDEP cases as compared to healthy participants, and patients (alive at the time of this analysis) classified as either at low or high risk of SUDEP based on the frequency of TCS (Experiment 2). Resting-state fMRI scans of 20 minute duration from each subject were considered for connectivity assessment in the ‘low GSA state’ and ‘high GSA state’. Seed-based connectivity analysis was employed focusing on twelve brain structures with a key role in autonomic and respiratory regulation. The seed-based correlation maps were further analyzed with PCA to summarize differences in connectivity between the groups and states in terms of a few components that explain most of the variance in the data.

Consistent with previous studies (Chang and Glover, 2009; Deen et al., 2011; Seeley et al., 2007; Uddin et al., 2009), activity in the anterior insula was found to be positively correlated with that in the anterior cingulate cortex and inferior parietal lobe (‘anterior insula positive network’), and negatively correlated with PCu/PCC and lateral parietal cortices (‘anterior insula negative network’; Fig. 4A). However, our analysis yielded an FC pattern (in the form of the 3^rd^ principal component) with a spatial map that is, to a large extent, the inverse of the mean anterior insula connectivity (Fig. 4B) and whose involvement in epilepsy patients was relatively high (Fig. 4C-Fig. 4D). This finding indicates weaker anterior insula networks in epilepsy patients compared to healthy controls. Furthermore, this weakening was more pronounced in the low state (characterized by more regular cardiac and breathing activity).

Our results also showed that the connectivity of cuneus and PCu/PCC is altered in epilepsy patients (low- and high-risk patients and SUDEP cases; Suppl. Fig. 3-Suppl. Fig. 4), which is in line with previous findings (Kay et al., 2013; Luo et al., 2012; Rajpoot et al., 2015); However, we found no evidence specifically implicating the aforementioned regions in SUDEP. Namely, the principal components that discriminated epilepsy patients from healthy controls did not reveal significant differences between SUDEP cases and low- or high-risk patients.

### 4.1 Relationship of anterior insula connectivity to survival time from fMRI scan

The observation of a negative correlation between the strength of anterior insula connectivity during the high state (irregular cardiorespiratory activity) and the interval between the fMRI scan and time of SUDEP may point to a predictive marker. Specifically, we found that patients who died sooner after the fMRI scan (2-4 years post-scan) exhibited lower anterior insula connectivity compared to patients who died later (5-8 years post-scan; *p* = 0.02, uncorrected; N = 9; Fig. 5). While this suggests an important dysfunctional role specific to the insula in SUDEP, it should be pointed out that the strength of anterior insula connectivity with its associated positive and negative networks was similar in SUDEP cases to that for low- and high-risk patients. Therefore, it is likely that other factors are required to coexist with weak connectivity in anterior insula to predispose individuals with epilepsy to SUDEP.

The anterior insula is known to play a major role in cardiovascular and respiratory functions, and has reciprocal connections with several limbic structures, including the anterior cingulate cortex, amygdala and hypothalamus (Palma and Benarroch, 2014). It receives viscerosensory inputs and, through its projections to brainstem output nuclei, contributes to regulation of blood pressure and other autonomic responses (Benarroch, 1993; Oppenheimer and Cechetto, 2016; Palma and Benarroch, 2014; Ruggiero et al., 1987; Saper, 1982). Functional imaging studies consistently report activation of the anterior insula in a wide range of interoceptive stimuli including dyspnea, ‘air hunger’ and heartbeat awareness, as well as in emotional processing (Brannan et al., 2001; Craig, 2009; Harrison et al., 2021; Zaki et al., 2012). In epilepsy, it has been suggested that seizures may affect insula activity leading to respiratory depression and cardiac arrhythmia, thereby increasing the risk of SUDEP (Li et al., 2017; Oppenheimer, 2001). However, scalp EEG, as used typically in the clinic for detecting seizures, has poor sensitivity to electrical activity from deep structures such as the insula and, as a consequence, it is difficult to assess whether seizure-induced autonomic manifestations are caused by insular dysfunction (Oppenheimer and Cechetto, 2016).

Altered FC between the insula and cingulate cortex in SUDEP and high-risk cases has been previously shown (Allen et al., 2019a), which is in line with our findings. In addition, patients that died of SUDEP at a time close to a structural MRI scan exhibited increased volume of the anterior insula (Allen et al., 2019b), further supporting the notion that impairment in the anterior insula may contribute to SUDEP. Finally, we recently found, in epilepsy patients, abnormal connectivity of the insula with the thalamus relative to changes in cardiac rhythms (Kassinopoulos et al., 2021). Collectively, these findings suggest that, in epilepsy, the communication of the anterior insula with other regions of the central nervous system may deteriorate over time, potentially leading to increased risk of cardiorespiratory failure.

### 4.2 Impaired communication between ventral and default mode network in epilepsy patients

We observed that activity in the anterior insula is positively correlated with that in regions of the ventral network and negatively correlated with regions of the default mode network (Fig. 4), and that this effect is weaker in patients with epilepsy, and is linked to SUDEP (Fig. 5). Seeley et al. (2007) first identified the so-called ventral network (also known as salience network) consisting of the anterior insula and anterior cingulate cortex as well as subcortical and limbic structures, with their activity linked to measures of anxiety. Subsequent functional studies implicated ventral attention regions mediating sympathetic activity (Beissner et al., 2013). Similarly, regions of the default mode network, including the posterior cingulate cortex, were found to be associated with parasympathetic activity (Beissner et al., 2013). Sridharan et al. (2008) demonstrated that the salience network, and particularly the anterior insula, is responsible for reducing activity in the default mode network during goal-directed tasks. Initial studies attributed autonomic dysfunction in epilepsy to seizures originating from, or spreading to, individual regions such as the anterior insula (Oppenheimer, 2001) and anterior cingulate cortex (Devinsky et al., 1995). However, our results suggest also the possibility that cardiorespiratory failure may rise from impaired communication between the ventral and default mode network due to insula dysfunction which could also explain the alteration in parasympathetic and sympathetic activity observed often in individuals with epilepsy (Myers et al., 2018).

### 4.3 Limitations and possible future work

Our study has a number of limitations: The unavailability of concurrent PPG and respiratory belt recordings led us in using GSA as a probe of cardiac and breathing activity. Although GSA reflects physiological changes, it cannot distinguish between irregular cardiovascular (transient increase in heart rate, variations in PPG amplitude, variations in blood pressure) and breathing (transient apnea, variations in tidal volume, etc.), with possible loss of sensitivity. For most witnessed SUDEP events, a TCS is observed a few minutes before death which triggers abnormal breathing patterns followed by episodes of bradycardia and terminal asystole, pointing to possible future studies focused on the link between irregular breathing and brain connectivity.

Our data was collected during daytime. SUDEP often occurs at night and likely during sleep when physiological processes change (e.g. heart and respiratory rates initially decline, but then increase and become more variable on entering rapid eye movement sleep; Purnell et al., 2018). Given these changes, in the present study, we sought to assess FC at different physiological states. However, the increased risk of SUDEP at night may result from other factors such as the influence of circadian rhythms and sleep on the brain, which could not be studied here since the fMRI data had been collected during daytime in a resting condition. While studying FC during sleep could help us understand the mechanisms underlying cardiorespiratory dysfunction, such studies are rare, as sleeping inside the scanner overnight is not well tolerated by participants. Therefore, fMRI studies need to consider protocols which mimic conditions that likely contribute to cardiorespiratory failure, while also being safe and practical. An example of such a test is the hypercapnic ventilatory response which was shown by Sainju et al. (2019) to indicate individuals with a prolonged increase in post-ictal CO_2_ after TCS, that are arguably at a high risk of SUDEP. Future fMRI studies may need to consider breathing or cardiovascular challenges during scans, along with physiological monitoring, to better understand how regional brain areas respond to such manipulation in patients at risk for SUDEP.

Finally, although our results suggest that the strength of anterior insula connectivity is inversely proportional to the interval between the fMRI scan and time of SUDEP (*r* = -0.74; *p* = 0.02, uncorrected; N = 9; Fig. 5), this finding is based solely on cross-sectional data and, thus, should be interpreted with caution. Longitudinal data are needed to directly address whether insular functions deteriorate over time in patients that later die of SUDEP or whether measures of anterior insula connectivity can be used as biomarkers of SUDEP risk.

## 5. Conclusions

In this work we revealed altered FC patterns of cuneus and precuneus/posterior cingulate cortex in epilepsy as compared to healthy controls during periods of regular cardiorespiratory rhythms. This was based on the use of global signal amplitude (GSA), which we showed to be a surrogate of cardiorespiratory rhythm irregularity (transient apnea, transient increase in heart rate, etc.). In addition, we found reduced anterior insula connectivity in epilepsy, particularly during regular cardiac and breathing activity. For SUDEP cases, this effect was inversely correlated with the scan-SUDEP interval. Overall, our results suggest that connectivity measures of anterior insula are potential biomarkers for SUDEP risk. Future research efforts should focus on gaining insight into the role of anterior insula connectivity in seizure-induced cardiorespiratory failure and how intervention strategies could be employed to restore anterior insula function.

## Acknowledgments

This work was funded by Epilepsy Research UK (Grant number: P1905). Funding by the NIH—National Institute of Neurological Disorders and Stroke (The Center for SUDEP Research; Grant number: U01-NS090407) and Hungarian NRDIO (National Research, Development, and Innovation Office; Grant number: OTKA K128040) is also acknowledged. Thanks to Luke Allen for help with data curation. We are grateful to the Epilepsy Society for supporting the Epilepsy Society MRI scanner. This work was supported by the National Institute for Health Research University College London Hospitals Biomedical Research Centre.

## Supplementary Material

**Suppl. Table 1.**
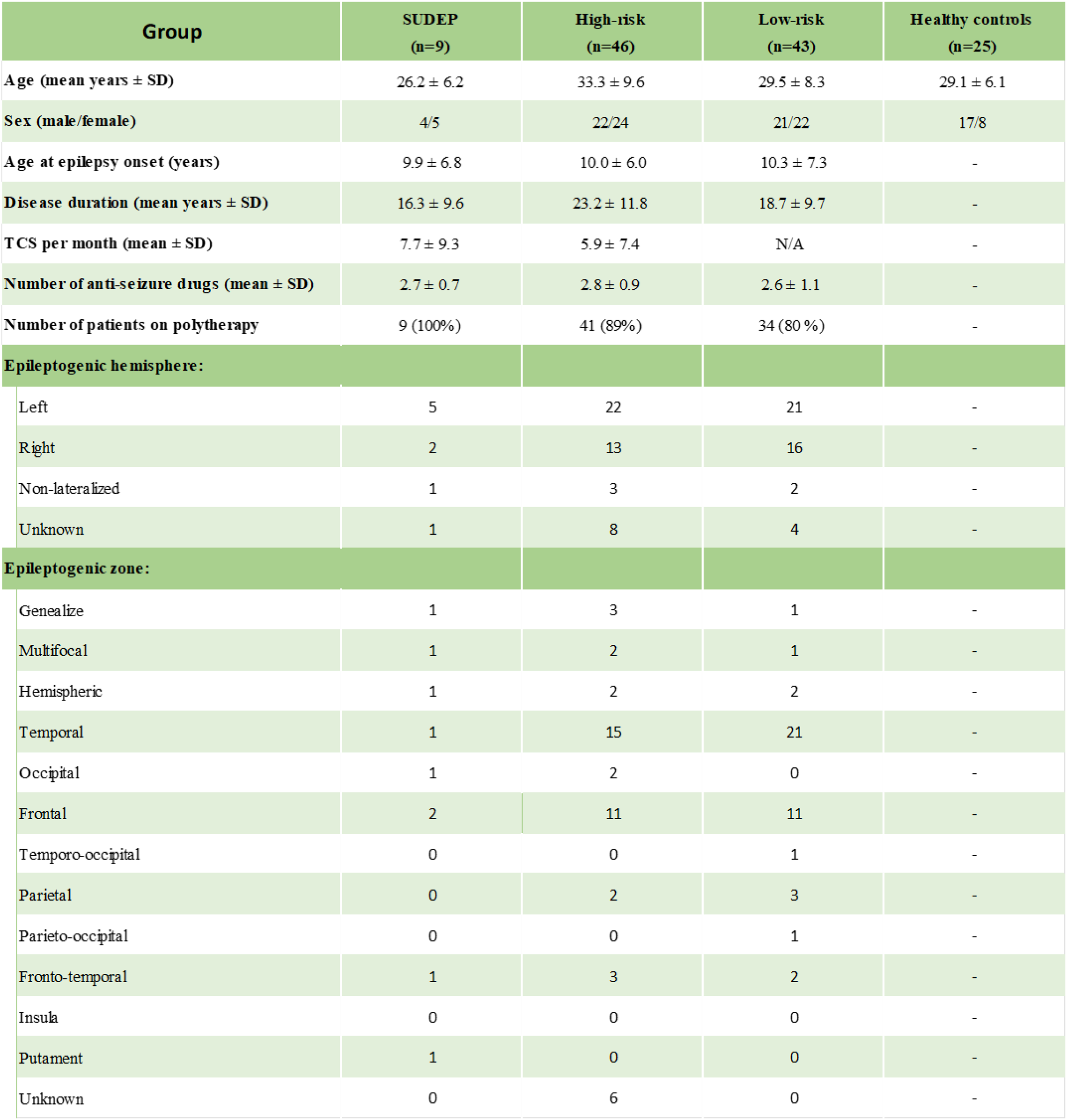
Exp. 1. Group demographic and clinical summaries of epilepsy patients and healthy controls. SD=standard deviation, N.A.=not applicable, TCS=tonic-clonic seizures.s

## Supplementary Methods

### a) Experiment 1. Preprocessing of the Human Connectome Project (HCP) dataset

We used resting-state fMRI scans from the HCP S1200 release (Van Essen et al., 2013) to examine the association of global signal amplitude (GSA) with variations in physiological variables. The HCP dataset includes, among others, T1-weighted (T1w) images and resting-state fMRI data (eyes-open and fixation on a cross-hair) from healthy young individuals (age range: 22-35 years) acquired on two different days. On each day, two 15-minute scans were collected (TR = 0.72 s). The preprocessing of the fMRI dataset is described in detail in Glasser et al. (2013). In the present study, the first scan from Day 1 was considered from 400 subjects who had good quality photoplethysmograph (PPG) and respiratory signal, as assessed by visual inspection.

The timings of the peaks in PPG were used to derive the heart rate, while the amplitudes of the PPG peaks were used to model the low-frequency variations in the envelope of the PPG signal. As described previously (Kassinopoulos and Mitsis, 2021, 2019), to facilitate peak detection, the PPG signal was initially band-pass filtered with a 2nd order Butterworth filter between 0.3 and 10 Hz. The minimum peak distance specified for peak detection varied between 0.5 and 0.9 s, depending on the subject’s average heart rate. The heart rate signal was computed in beats-per-minute (bpm) and evenly resampled at 10 Hz. The amplitudes of the peaks were also evenly resampled at 10 Hz. The resampling of heart rate and PPG-Amp was done using linear interpolation. The breathing signal was detrended linearly and corrected for outliers using a median filter. Subsequently, the breathing signal was low-pass filtered at 5 Hz with a 2nd order Butterworth filter and z-scored. To extract the fMRI global signal of each scan, we initially performed tissue segmentation on the T1w images in the MNI152 space using FLIRT in FSL 5.0.9, which generated probabilistic maps for the grey matter, white matter and cerebrospinal fluid compartments (Zhang et al., 2001). Afterwards, the global signal was calculated by estimating the mean time-series across all voxels with a probability of belonging to GM above 0.25.

### b) Experiment 2. Preprocessing of electrocardiogram (ECG)

The ECG was corrected for gradient artifacts using adaptive template subtraction (Allen et al., 2000) implemented in BrainVision Analyzer 2 software (Brain Products GmbH, Munich, Germany), and band-pass filtered from 0.5 to 40 Hz. The R-waves were detected using Matlab’s function *findpeaks* with a minimum peak distance varying between 0.5 and 0.9 s depending on the subject’s average RR interval (time between successive R-waves).

**Suppl. Fig. 1.**
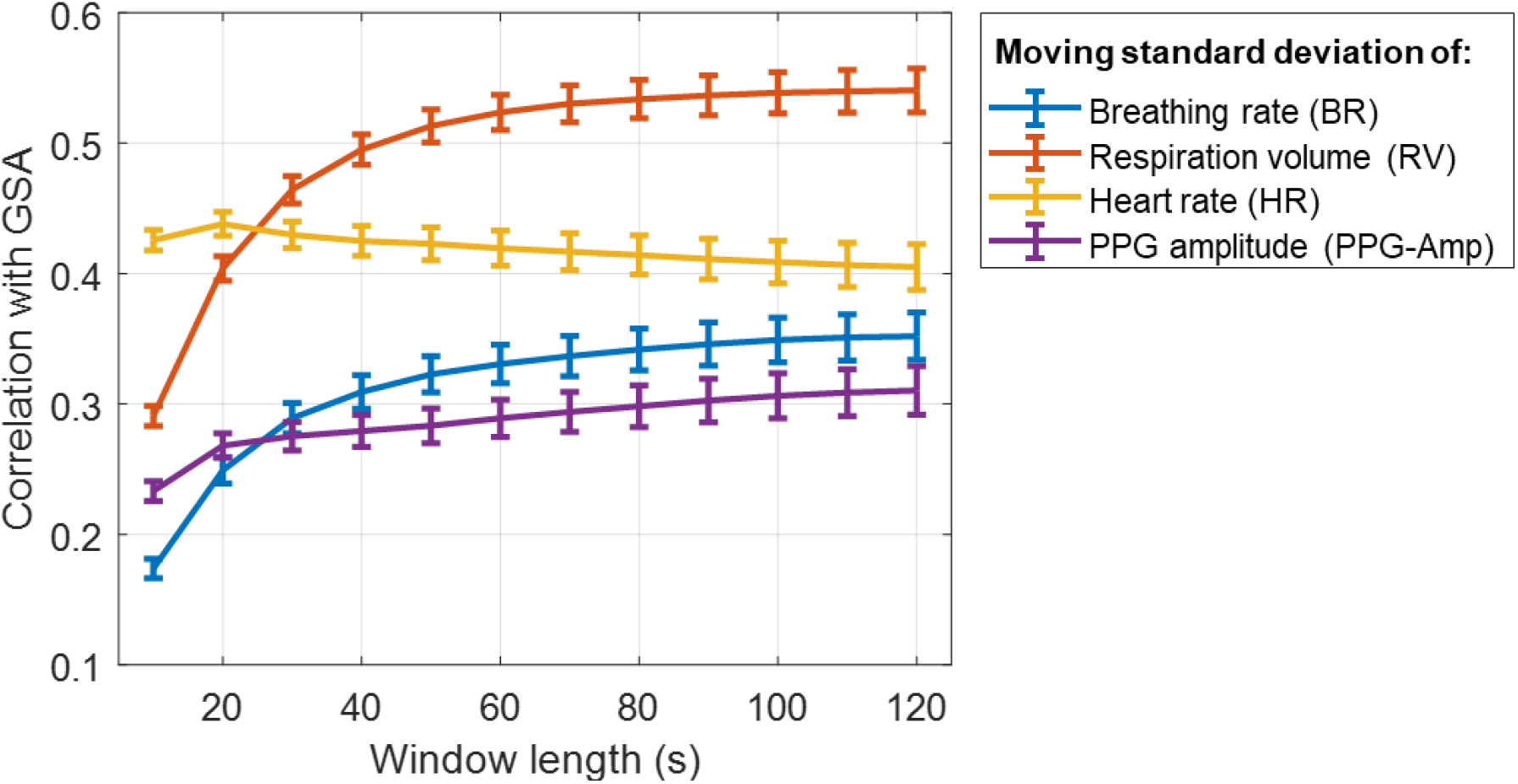
Exp. 1. Correlation of global signal amplitude (GSA) with the moving standard deviation of physiological variables for different window lengths. Resting-state fMRI data and concurrent photoplethysmograph (PPG) and breathing recordings from 400 healthy young subjects of the Human Connectome Project (Van Essen et al. 2013) were used for this analysis. The moving standard deviations of breathing rate, respiration volume, heart rate and PPG amplitude were computed for window lengths ranging from 10 to 120 sec. Similarly, GSA (i.e. moving standard deviation of fMRI global signal) was computed for the same window lengths, and its correlation with the traces of the physiological variables was estimated and averaged across 400 subjects (error bars indicate standard errors). Increasing the window length from 20 to 80 sec was found to enhance the correlation of GSA with the amplitude of variations in breathing rate, respiration volume and PPG amplitude, whereas no substantial additional benefit was observed for longer lengths.

**Suppl. Fig. 2.**
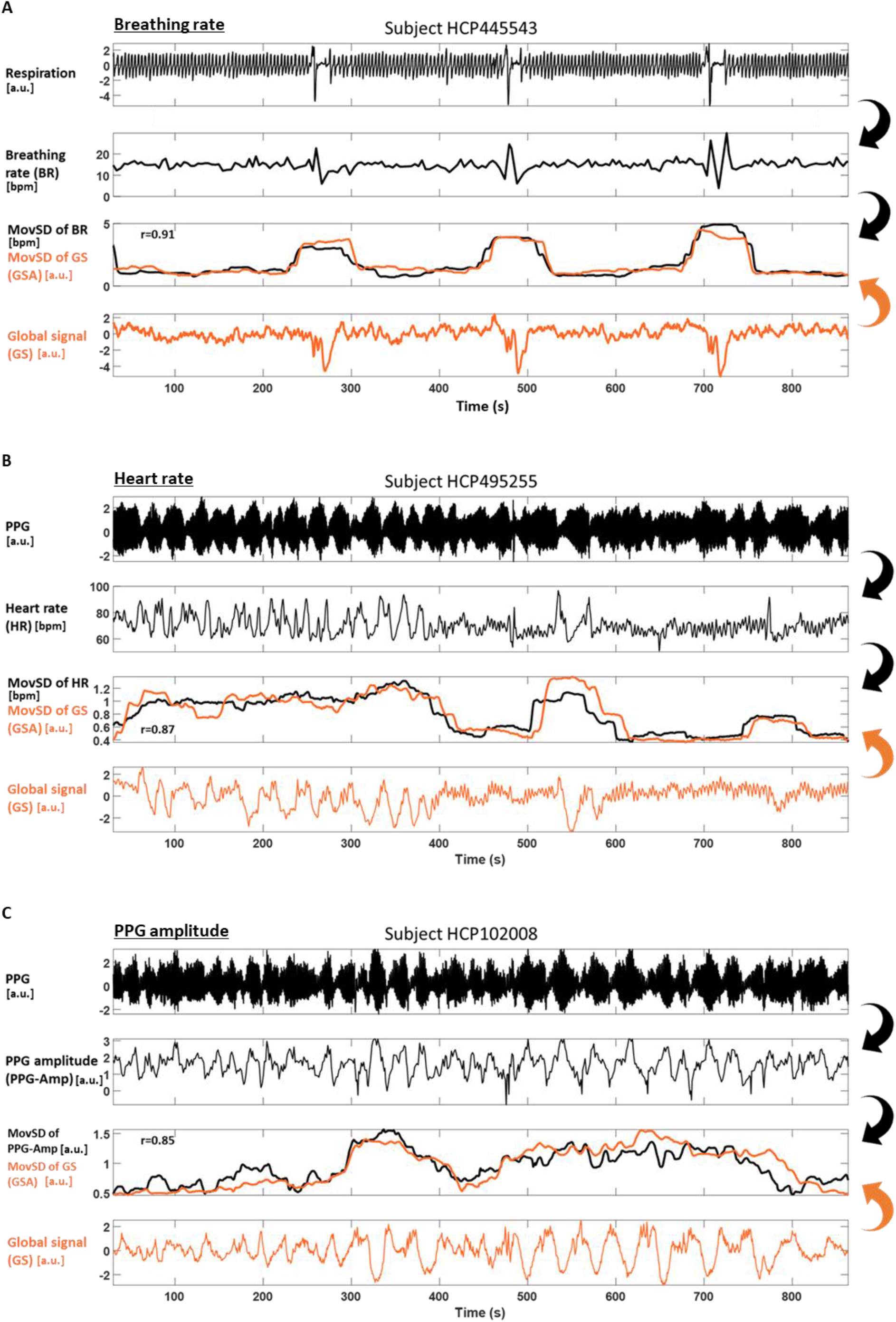
(Next page). Exp 1. Association of GSA with variations in (A) breathing rate, (B) heart rate and (C) PPG amplitude, during rest. In each panel, the first row shows the raw physiological signal (respiration or PPG), the second row shows the physiological variable extracted from the raw signal (i.e. breathing rate, heart rate or PPG amplitude), and the third row shows in black color the moving standard deviation of the physiological variables (window length: 80 sec). Overall, we observe that an increase in the levels of GSA can be induced by a transient apnea (HCP445535), a transient increase in heart rate (HCP495255) or strong fluctuations in PPG amplitude (HCP102008).

**Suppl. Fig. 3.**
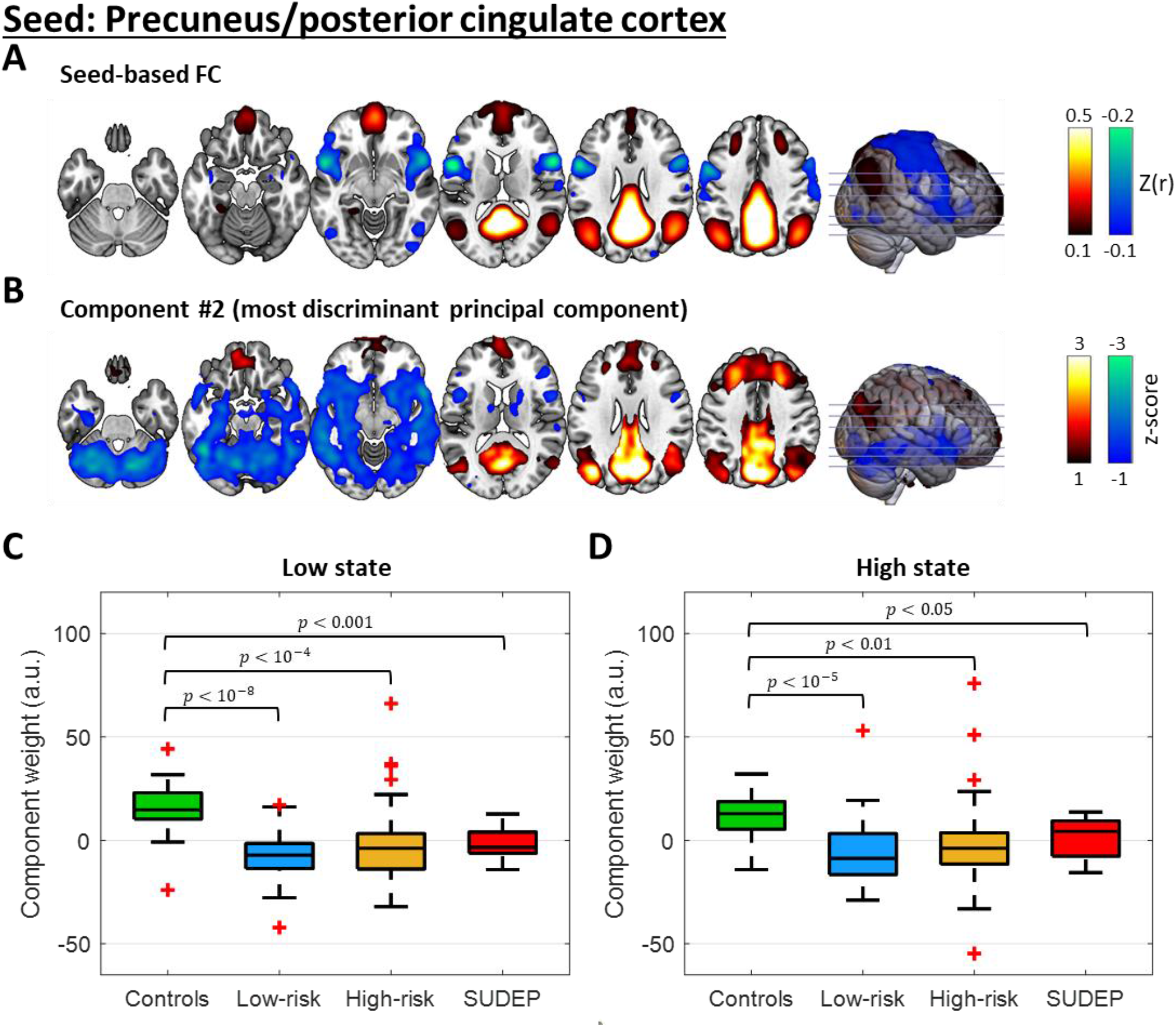
Exp. 2. Involvement of FC pattern #2 of precuneus/posterior cingulate (PCu/PCC) connectivity in the low and high state. **(A)** Seed-based correlation map with the seed placed in the PCu/PCC averaged across subjects and time. **(B)** FC pattern of component #2 derived from the PCu/PCC connectivity profiles of all subjects through PCA. Component weights of the four groups in the **(C)** low and **(D)** high state. For the spatial involvement of Yeo large-scale networks of FC pattern #2, see Suppl. Fig. 5.

**Suppl. Fig. 4.**
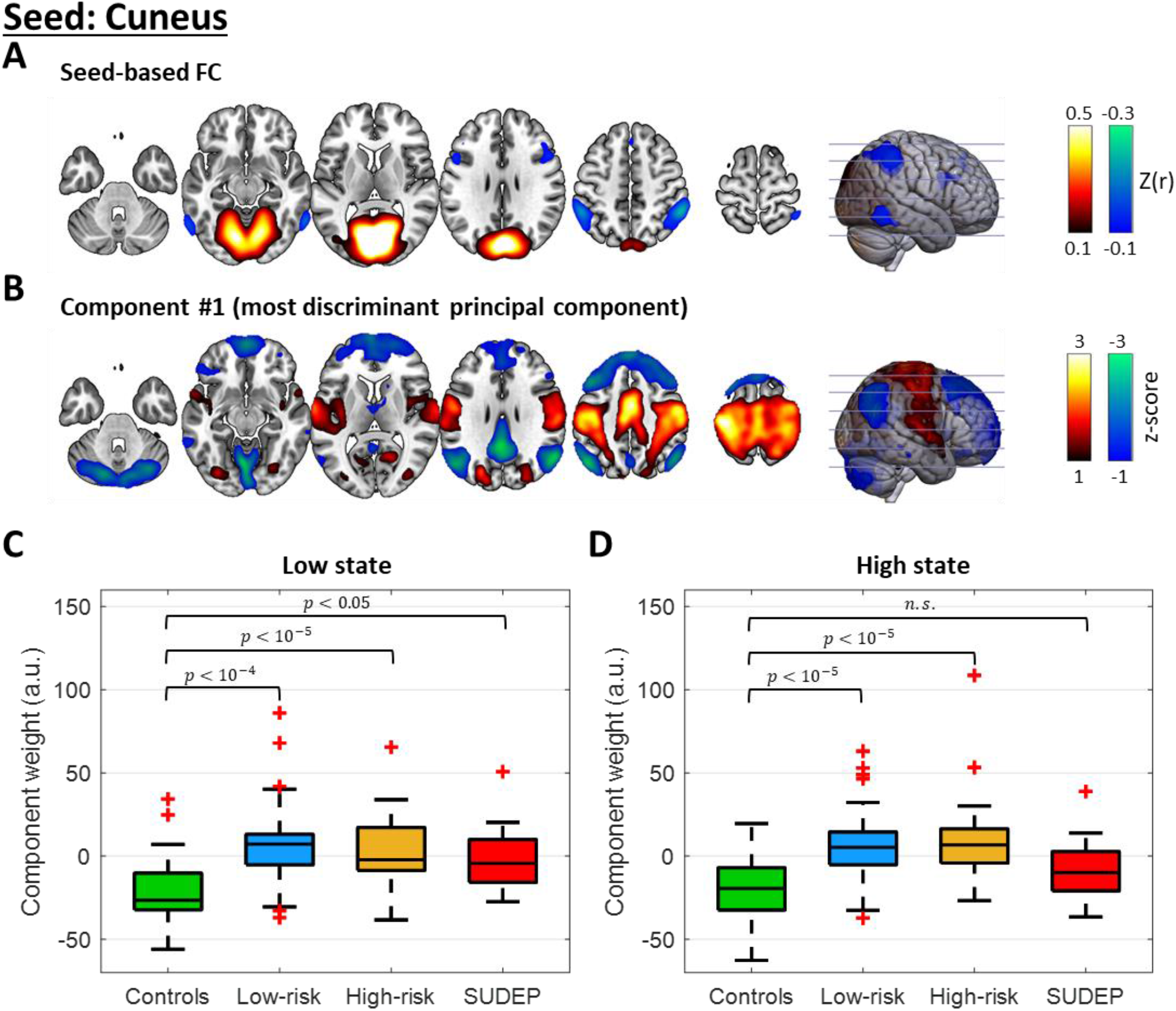
Exp. 2. Involvement of FC pattern #1 of cuneus in the low and high state. **(A)** Seed-based correlation map with the seed placed in the cuneus averaged across subjects and time. **(B)** FC pattern of component #1 from the cuneus connectivity profiles of all subjects using PCA. Component weights of the four groups in the **(C)** low and **(D)** high state. For the spatial involvement of the Yeo large-scale networks of FC pattern #1, see Suppl. Fig. 5.

**Suppl. Fig. 5.**
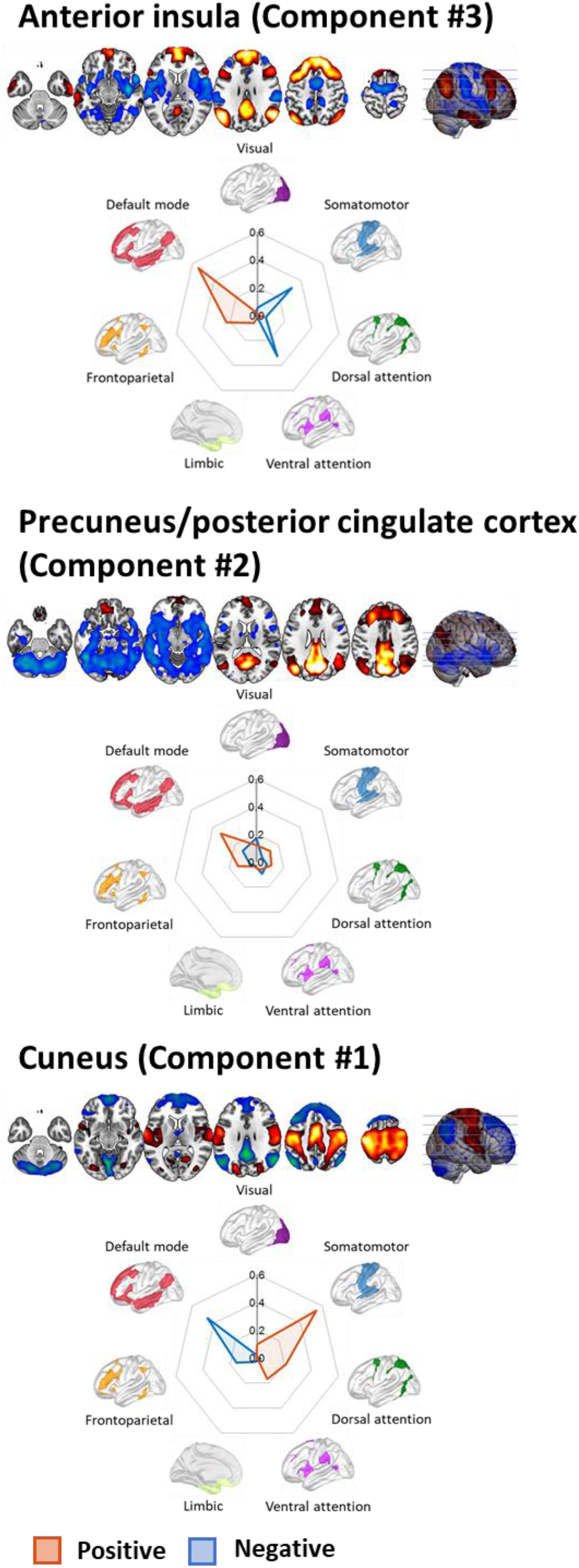
Exp. 2. Spatial involvement of the seven large-scale networks of the Yeo atlas (Yeo et al., 2011) in the FC patterns of the most discriminant components. The spatial involvement with the networks was assessed by calculating the Sørensen–Dice coefficient using the ICN_Atlas toolbox (Kozák et al., 2017) for the most discriminant components of the FC patterns, namely those that exhibited an *F*-statistic above chance level (*F* = 9.4, *p* < 10^-4^; Fig. 3) for the anterior insula (top), PCu/PCC (middle) and cuneus (bottom). The sign of the components’ constituent regions is represented in the involvement plots as red for positive and blue for negative.

