## Supplementary Material for "Brain Connectivity Correlates of Breathing and Cardiac Irregularities in SUDEP: A Resting-State fMRI Study"

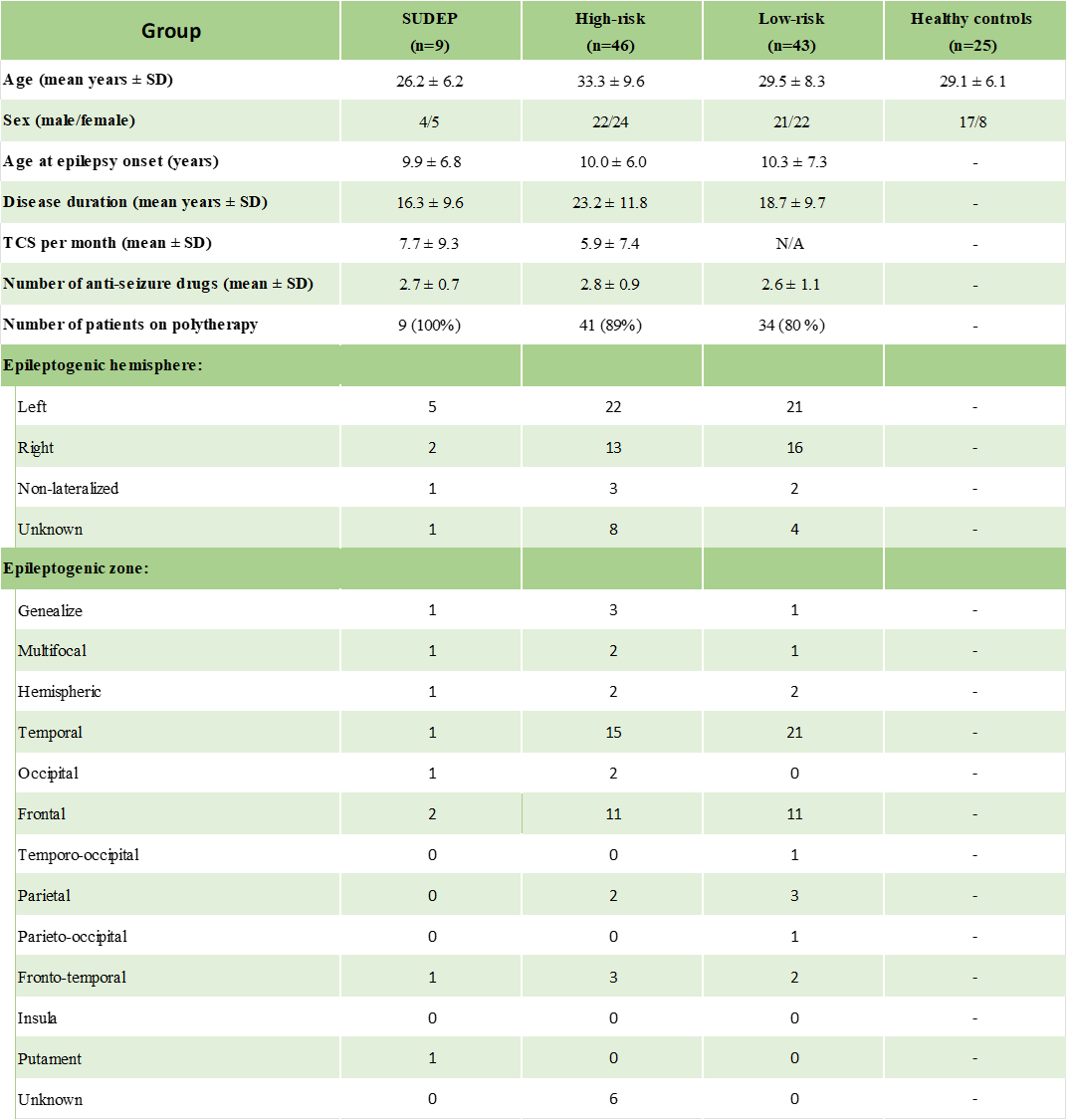

**Suppl. Table 1. Exp. 1. Group demographic and clinical summaries of epilepsy patients and healthy controls.** SD=standard deviation, N.A.=not applicable, TCS=tonic-clonic seizures.

**Supplementary Methods**

**Suppl. Fig. 1.** **Exp. 1. Correlation of global signal amplitude (GSA) with the moving standard deviation of physiological variables for different window lengths.** Resting-state fMRI data and concurrent photoplethysmograph (PPG) and breathing recordings from 400 healthy young subjects of the Human Connectome Project (Van Essen et al. 2013) were used for this analysis. The moving standard deviations of breathing rate, respiration volume, heart rate and PPG amplitude were computed for window lengths ranging from 10 to 120 sec. Similarly, GSA (i.e. moving standard deviation of fMRI global signal) was computed for the same window lengths, and its correlation with the traces of the physiological variables was estimated and averaged across 400 subjects (error bars indicate standard errors). Increasing the window length from 20 to 80 sec was found to enhance the correlation of GSA with the amplitude of variations in breathing rate, respiration volume and PPG amplitude, whereas no substantial additional benefit was observed for longer lengths.

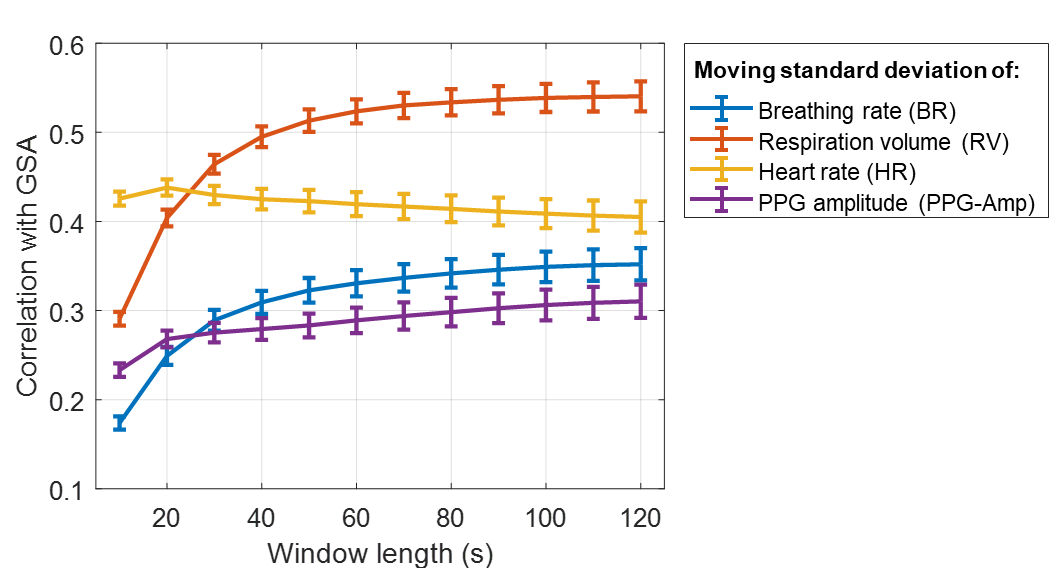

**Suppl. Fig. 2 (Next page). Exp 1. Association of GSA with variations in (A) breathing rate, (B) heart rate and (C) PPG amplitude, during rest.** In each panel, the first row shows the raw physiological signal (respiration or PPG), the second row shows the physiological variable extracted from the raw signal (i.e. breathing rate, heart rate or PPG amplitude), and the third row shows in black color the moving standard deviation of the physiological variables (window length: 80 sec). Overall, we observe that an increase in the levels of GSA can be induced by a transient apnea (HCP445535), a transient increase in heart rate (HCP495255) or strong fluctuations in PPG amplitude (HCP102008).

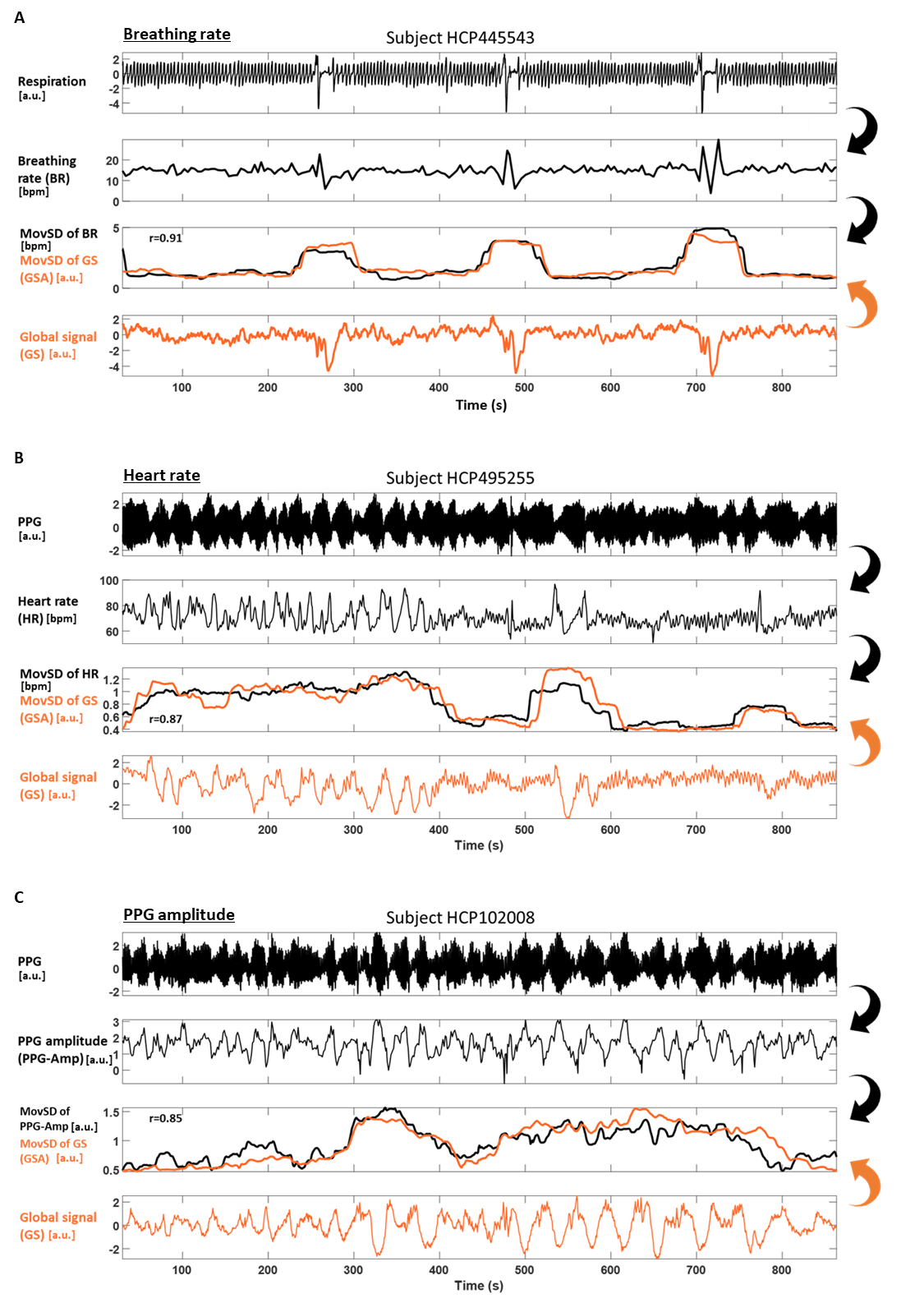

**Suppl. Fig. 3. Exp. 2. Involvement of FC pattern #2 of precuneus/posterior cingulate (PCu/PCC) connectivity in the low and high state. (A)** Seed-based correlation map with the seed placed in the PCu/PCC averaged across subjects and time. **(B)** FC pattern of component #2 derived from the PCu/PCC connectivity profiles of all subjects through PCA. Component weights of the four groups in the **(C)** low and **(D)** high state. For the spatial involvement of Yeo large-scale networks of FC pattern #2, see Suppl. Fig. 5.

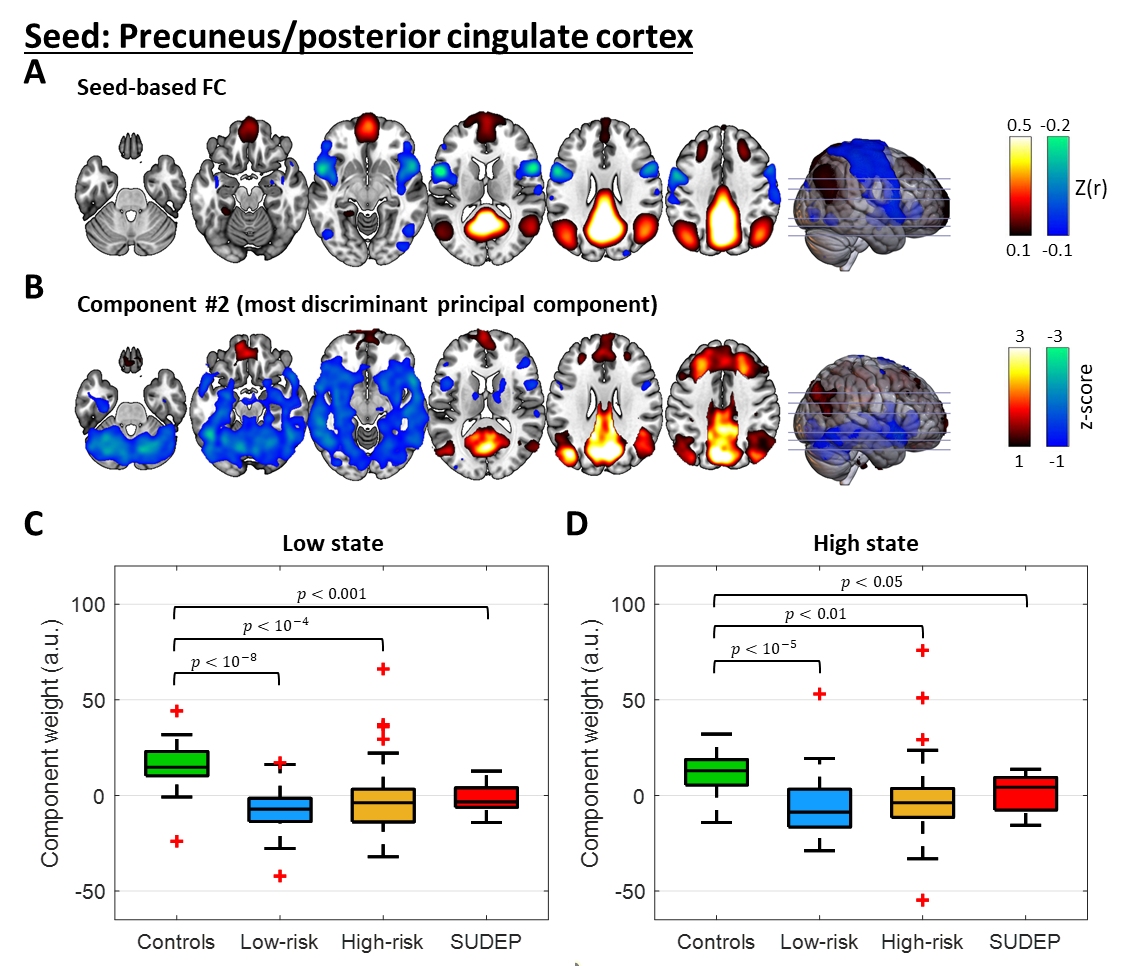

**Suppl. Fig. 4.** **Exp. 2. Involvement of FC pattern #1 of cuneus in the low and high state. (A)** Seed-based correlation map with the seed placed in the cuneus averaged across subjects and time. **(B)** FC pattern of component #1 from the cuneus connectivity profiles of all subjects using PCA. Component weights of the four groups in the **(C)** low and **(D)** high state. For the spatial involvement of the Yeo large-scale networks of FC pattern #1, see Suppl. Fig. 5.

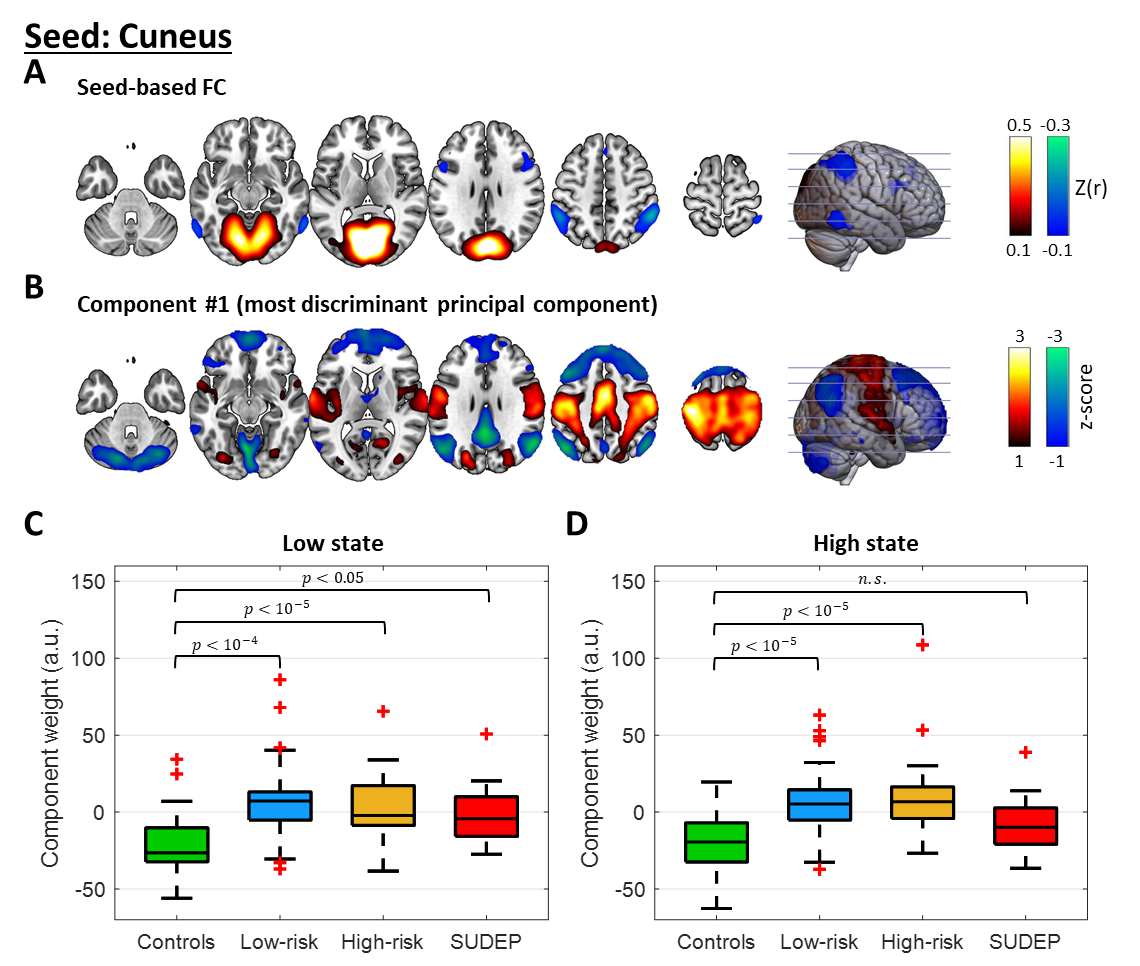

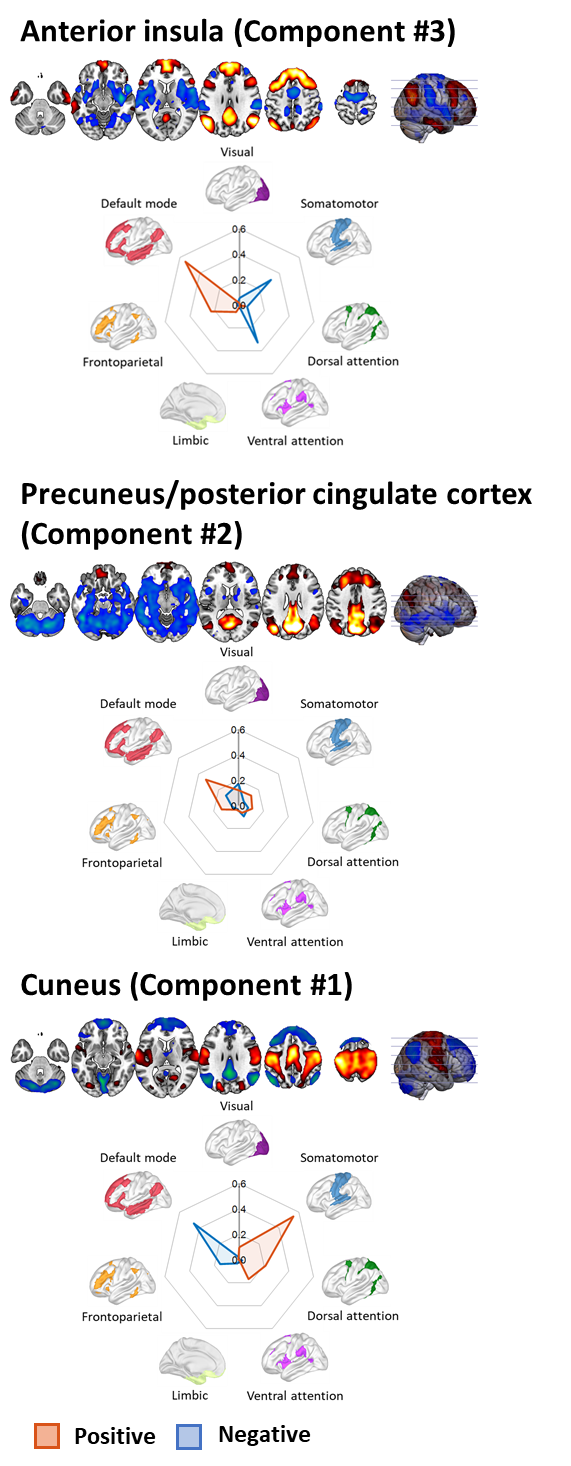

**Suppl. Fig. 5. Exp. 2. Spatial involvement of the seven large-scale networks of the Yeo atlas (Yeo et al., 2011) in the FC patterns of the most discriminant components.** The spatial involvement with the networks was assessed by calculating the Sørensen–Dice coefficient using the ICN_Atlas toolbox (Kozák et al., 2017) for the most discriminant components of the FC patterns, namely those that exhibited an *F*-statistic above chance level (*F* = 9.4, *p* < 10^-4^; Fig. 3) for the anterior insula (top), PCu/PCC (middle) and cuneus (bottom). The sign of the components’ constituent regions is represented in the involvement plots as red for positive and blue for negative.
